# Methods for making and observing model lipid droplets

**DOI:** 10.1101/2023.07.17.549385

**Authors:** Sonali A. Gandhi, Shahnaz Parveen, Munirah Alduhailan, Ramesh Tripathi, Nasser Junedi, Mohammad Saqallah, Matthew A. Sanders, Peter M. Hoffmann, Katherine Truex, James G. Granneman, Christopher V Kelly

**Affiliations:** Department of Physics and Astronomy, Wayne State University, Detroit, MI, USA 48201; Center for Molecular Medicine and Genetics, School of Medicine, Wayne State University, Detroit, MI, USA 40201; Physical Sciences Department, Embry-Riddle Aeronautical University, Daytona Beach, FL, USA 32114; Department of Physics, United States Naval Academy, Annapolis, MD, USA 21402; Center for Integrative Metabolic and Endocrine Research, School of Medicine, Wayne State University, Detroit, MI USA 48201

**Keywords:** Model lipid droplets, fluorescence microscopy, atomic force microscopy, fluorescence recovery after photobleaching, fluorescence correlation spectroscopy, fluorescence lifetime imaging microscopy, phospholipid monolayers

## Abstract

The mechanisms by which the lipid droplet (LD) membrane is remodeled in concert with the activation of lipolysis incorporate a complex interplay of proteins, phospholipids, and neutral lipids. Model LDs (mLDs) provide an isolated, purified system for testing the mechanisms by which the droplet composition, size, shape, and tension affects triglyceride metabolism. Described here are methods of making and testing mLDs ranging from 0.1 to 40 µm diameter with known composition. Methods are described for imaging mLDs with high-resolution microscopy during buffer exchanges for the measurement of membrane binding, diffusion, and tension via fluorescence correlation spectroscopy (FCS), fluorescence recovery after photobleaching (FRAP), fluorescence lifetime imaging microscopy (FLIM), atomic force microscopy (AFM), pendant droplet tensiometry, and imaging flow cytometry. These complementary, cross-validating methods of measuring LD membrane behavior reveal the interplay of biophysical processes in triglyceride metabolism.

## Introduction

Cellular homeostasis depends on a precise metabolic balance of sugars and lipids through times of consumption and fasting. The storage and use of oil-based energy such as triacylglycerols and sterol esters in lipid droplets (LDs) is a nearly ubiquitous cellular means of maintaining healthy free fatty acid levels.^1–3^ Most cells maintain LDs between 0.1 and 100 µm diameter with an oily core surrounded by a monolayer of phospholipids (PLs) and proteins depending on the cell type.^4–6^ LDs grow between the leaflets of the endoplasmic reticulum (ER) bilayer with proteins spontaneously sorting between the ER bilayer and the LD monolayer.^7, 8^ In addition to maintaining energy balance, lipid droplets also facilitate intracellular coordination between organelles^9–11^ and are engineered for pharmaceutical production and delivery.^12, 13^

Cells carefully regulate the LDs composition, size, and number per cell, but the molecular and biophysical mechanisms of LD nucleation and growth remain unknown. Properties such as the LD membrane curvature and tension affect the LD budding directionality and fission from the ER,^14, 15^ but variables such as the LD composition, neutral lipid production rates, and LD-nucleating proteins have unresolved importance. Intracellular trafficking balances protein concentrations on the LD monolayer, on membrane bilayers, and in the cytosol.^16^ Perilipin 5 (PLIN5) is an LD-targeting and LD-regulating protein that has shown to have diverse importance in fatty acid storage and mitochondrial energy utilization.^17^ PLIN5 has many dynamic lipid and protein binding partners that depend on the cellular metabolic state, which includes a ligand-dependent unbinding from α/β hydrolase domain-containing protein 5 that activates lipolysis.^18, 19^ Protein crowding and competition for the limited LD monolayer area has been proposed as key for determining the LD monolayer composition^20^ with the specifics of the PL composition, tension, and curvature likely influencing the metabolism of triacylglycerols (TAGs).^21^ Simplified model systems are needed for determining the underlying biophysical properties and molecular contributions that govern LD behavior.

Model LDs (mLDs) were created through a variety of means for diverse purposes. The binding of small PL vesicles to TAG droplets forms a surrounding monolayer, but saturation of the monolayer with PL from small vesicles takes >20 min. Equilibrium may be reached faster by sonicating PL-TAG mixtures to create PL-coated, oil-filled mLDs when the volume of aqueous buffer exceeds the volume of the neutral lipid. The PL-to-TAG ratio influences the final mLD size consistent with determining the surface area-to-volume ratio. However, monolayers are most robustly created when either the PLs are dissolved within the neutral lipid prior to its exposure to the aqueous buffer or neutral lipids are incorporated between the PL leaflets of giant unilamellar vesicles (GUVs) to create droplet-embedded vesicles (DEVs).^21–24^ The buoyancy of oil-filled mLDs in an aqueous buffer complicates the imaging of mLDs or DEVs by inverted microscopy. Modifying the mLD composition^25^ or using confined imaging chambers addresses this challenges,^26^ but with new difficulties introduced.

This manuscript focuses on mLD fabrication and measurement methods toward testing the binding rate, surface diffusion, membrane targeting, and membrane tension changes dependent on the LD-bound proteins and PLs. We have identified and addressed the challenge of observing curved, buoyant mLDs with conventional high-resolution inverted microscopy methods. DEVs, for example, provide the ability to observe protein sorting between PL bilayers and mLD monolayers despite challenges in sample heterogeneity, high-resolution imaging, and quantitative analyses. Through the confining of micron-sized mLDs within microbead-supported glass chambers, the bottom of the buoyant mLDs could be observed without refractive distortion by the mLD surface or the disruptive proximity of a substrate; this method was used with fluorescence correlation spectroscopy (FCS) to measure the concentration-dependent diffusion of PLIN5 on mLDs. With nanoscale mLDs that were diffusing and dispersed throughout the buffer, fluorescence lifetime imaging microscopy (FLIM) was employed to measure the increase in PL packing caused by PLIN5 on the mLD membrane. Coverslip-bound mLD caps enabled high-resolution imaging of mLD membrane with short-working distance, oil-immersion objectives while maintaining accessibility of the surrounding aqueous buffer. Coverslip-bound mLD caps were used to measure the membrane binding kinetics, perform fluorescence recovery after photobleaching (FRAP), and measure the mLD surface tension with atomic force microscopy (AFM). The tension measurements by AFM were validated by a custom, cost-effective pendant droplet tensiometer. The combination of methods developed for making and observing mLDs enables new experimental means of measuring the biophysical properties that govern LD biology. The methods reported here were used to measure the binding rate of PL vesicles to mLDs, the monolayer-vs-bilayer sorting of PLIN5, the membrane-bound diffusion of PLIN5, and the effects of surfactants on the mLD tension. These mLD preparation and observation methods enable resolving the fundamental biophysics that govern LD biology.

## Results

### Droplet embedded vesicles (DEVs)

By mixing GUVs with droplets of TAG, TAG was incorporated between the PL leaflets to create DEVs with distinct PL monolayers and PL bilayers (Fig. 1). Brightfield microscopy showed the index of refraction contrast by the GUV and by the embedded mLD (Fig. 1B). Confocal fluorescence imaging enabled isolating distinct imaging planes with the buoyant mLD typically floating to the top of the GUV (Fig. 1C-F). The net buoyancy of the DEV depended on the volumes and densities of the aqueous buffer, the GUV interior, and the attached mLD. The relative density of the GUV interior and the surrounding buffer is controllable via the salt density and composition present during GUV formation and subsequent dilution, respectively. By using large molecular weight solutes (*i.e*., sucrose) to fill the GUVs during their formation, the GUVs and DEVs typically sink when diluted in physiological buffers.

**Figure 1.**
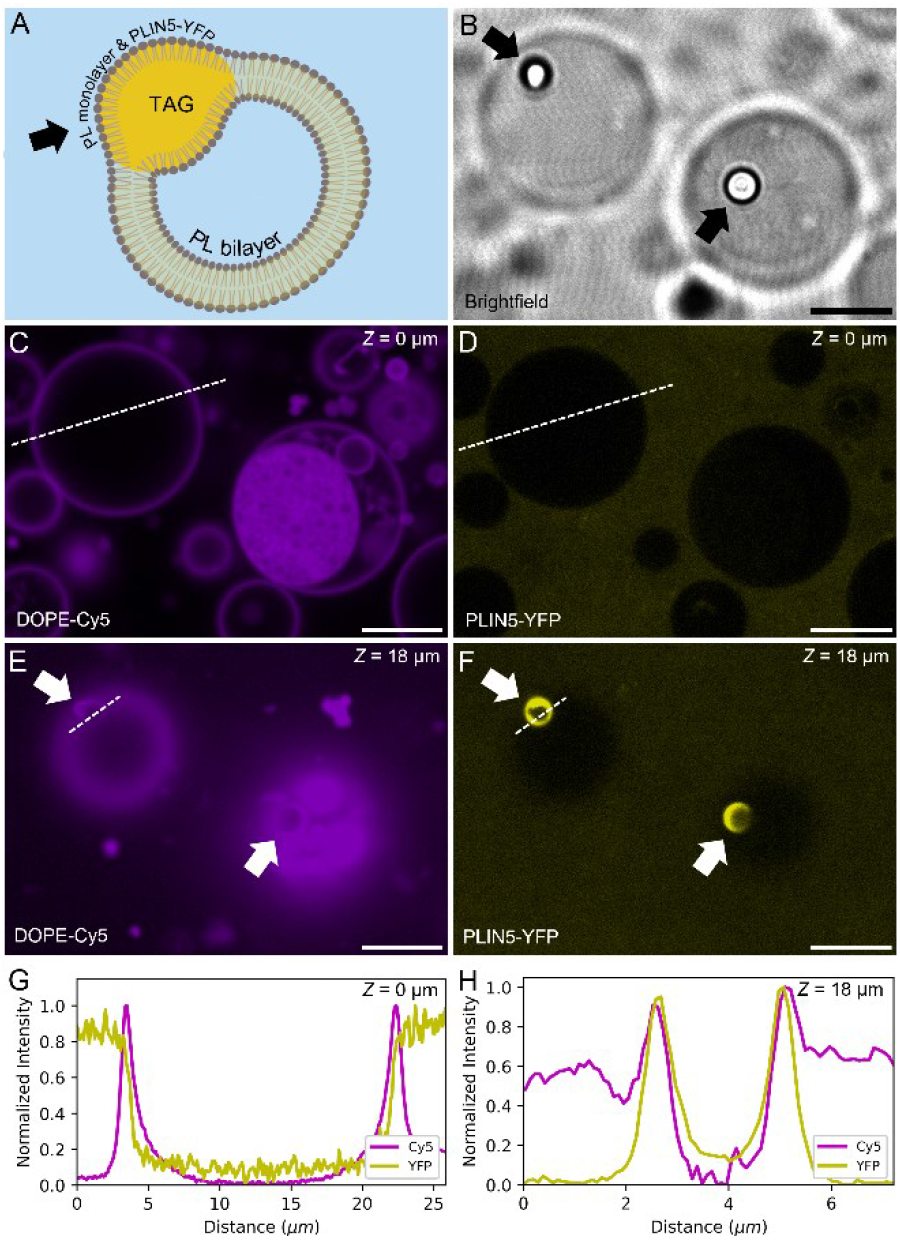
Droplet embedded vesicles (DEVs) (A) DEVs were made by adding TAG to GUVs. The oil sequesters between the leaflets of the PL bilayers to make mLDs attached to PL vesicles (*arrows*). (B) Brightfield microscopy showed a slight refractive index contrast between the GUVs and the surrounding buffer and a large index contrast between the mLDs the rest of the sample (*arrows*). (C and D) When focused on the center of the GUVs (*z* = 0), confocal microscopy showed the PL bilayers of the GUVs, the heterogeneity of the GUV sizes, and the vesicles within vesicles via the Cy5 channel (C). In the YFP channel (D), the 150nM of PLIN5-YFP in the surrounding buffer is present without any apparent accumulation on the PL bilayers. Scale bars represent 10 μm. (E and F) When focused on the center of the attached mLDs, the Cy5 channel (E) showed both the PL bilayers of the vesicles and the PL monolayers surrounding the mLDs. The YFP channel (F) showed the dense accumulation of PLIN5-YFP on the PL monolayer. Scale bars represent 10 μm. (G and H) Line scans through the two focal planes (*dotted lines*) show (G) no accumulation of PLIN5-YFP on the PL bilayer at *z* = 0 μm and (F) intense accumulation of PLIN5-YFP on the PL monolayer at *z* = 18 μm.

Quantifying the relative brightness in fluorescence imaging is complicated by the differential photobleaching of present fluorophores, the relative angle of the membranes to the imaging focal plane, and distortions caused by the curved surfaces of varying index of refraction. Electroformation generates large GUVs, but heterogeneity within the sample includes variation in the GUV sizes and nesting of vesicles within vesicles. When there are vesicles within vesicles, it becomes difficult to ensure that only one bilayer is contributing to the brightness of a given image pixel. Additionally, a membrane that is parallel to the microscopy focal plane (*i.e*., when focused on the top or bottom of the GUV) will yield dimmer pixel values than a membrane that is perpendicular (*i.e*., when focused on the equator of the GUV). The coupling of these effects can complicate the quantification of protein sorting even when the PL bilayer and the PL bilayer are both visible with a single image.

Confocal fluorescence imaging of DEVs effectively demonstrates the relative sorting of membrane-binding proteins between PL monolayers and PL bilayers despite the challenges inherent to DEVs.^21–24^ Shown here is the strong preference for PLIN5-YFP to bind to PL monolayers versus PL bilayers (Fig. 1). In the presence of 190 nM of PLIN5-YFP, confocal imaging showed PLIN5 in surrounding solution, no PLIN5 within the DEVs, PLIN5 bound and concentrated to the mLD surface, and no PLIN5 on the GUV bilayer. Line scans of fluorescence intensity show no correlated peaks in PLIN5-YFP with the PL bilayer when focused through the GUV (Fig. 1G); however, intense PLIN5-YFP emission was observed correlated with the PL when focused on the monolayer of the mLD (Fig. 1H).

### Spherical model lipid droplets (mLDs)

Spherical oil-in-water and water-in-oil droplets were created through sonication and vortexing mixtures of TAG, PL, and aqueous buffer. The successful creation of suspended droplets was apparent through a uniform cloudiness of the sample. Sonication proved to create independent, suspended droplets more reliably than vortexing, but a wide variability of sonication times was necessary depending on the specific bath sonicator used (*i.e*., between 10 sec and 2 min). Excessive sonication or vortexing resulted in a creaming of the mixture that was apparent as an oil-rich phase separation that was white and highly viscous. Creaming reduced the number and quality of the suspended mLDs available for further analysis.

The size of the oil-in-water droplets was affected by the ratio of TAG-to-PL; more TAG resulted in larger volume droplets whereas more PL resulted in more stabilization of the droplet surface area within the sample. When the volume of buffer far exceeded the volume of TAG, oil-in-water droplets were created, which suffer from droplets floating out of the field of view of inverted microscopes. When the volume of TAG exceeded the volume of buffer, water-in-oil droplets were created, which suffer from an inability expose the monolayer to additional aqueous components or altering the monolayer composition. Storing droplets on a rotator allows them to equilibrate and avoid coalescence prior to further analysis. To address the buoyancy complication of oil-in-water droplets during imaging, we created glass microscopy chambers with the mLDs stabilized between a microscopy coverslip and glass slide spaced with 20-µm diameter microspheres (Fig. 2A). Epifluorescence observation shows the fluorescent monolayer of the mLDs most clearly when focused on to the droplet’s equator and the monolayer extends into the optical z-axis (Fig. 2B, C).

**Figure 2.**
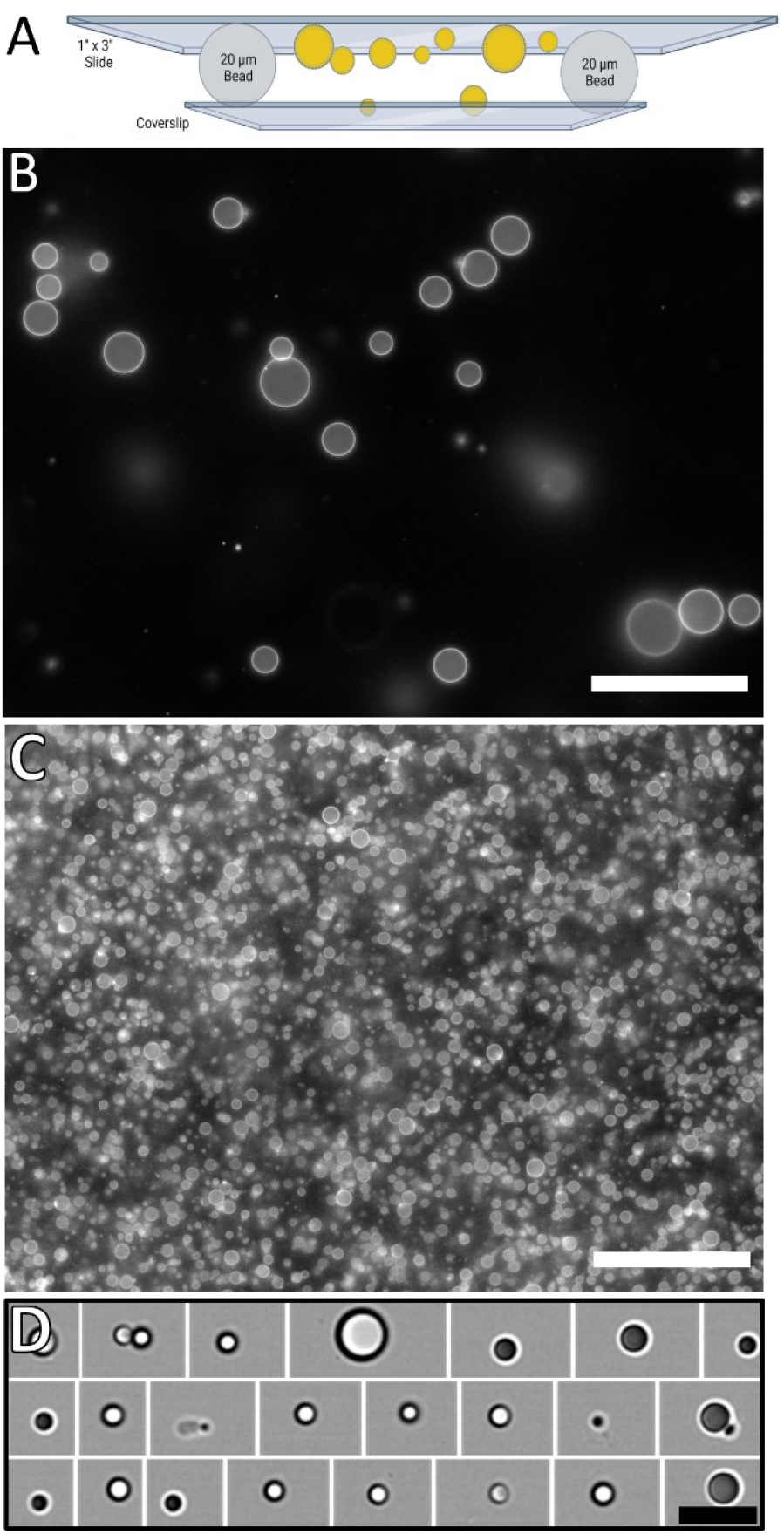
Buoyant oil-in-water mLDs. (A) Trapping the mLDs between a coverslip and a glass slide keeps the mLDs within the short working distance of oil immersion objectives. (B and C) Microscopy chambers were used to observe mLDs with (B) 1000 TAG:1 PL and (C) 100 TAG: 1 PL. Changing the z-focus of the objective allows discrimination of the equatorial plane from the glass surfaces in the imaging chamber (Fig. S1). (D) Alternatively, the oil-in-water mLDs were imaged in high-throughput with imaging flow cytometry, but without sequential imaging of any individual droplet. (B-D) Scale bars represent 40 µm.

Oil-in-water mLDs were also analyzed via imaging flow cytometry (Fig. 2D). The droplets were created by sonication in a microtube and introduced into the cytometer through the standard micro-tubing. The internal fluidics of the cytometer presented the droplets into the optical path for imaging with brightfield and fluorescence. The cytometer identified when a droplet was in view and saved the associated images. Images of over 100k droplets were acquired within 3 min and available for off-line analysis. Attempts to analyze mLD shape changes and protein sorting with imaging flow cytometry were complicated by protein-induced aggregation of the mLDs during the sample loading. Smaller mLDs bound together and were indistinguishable from larger mLDs changing shape.

### Stability and buffer exchange for mLD caps

A glass microscopy chamber was developed for imaging buoyant mLDs with inverted, high-numerical aperture objectives, but it prohibits the addition or exchange of the surrounding buffer. This limitation prohibited the resolution of time-dependent compositional changes to the monolayer, such as directly monitoring the binding rates of LD-binding PLs or proteins. To overcome this, we developed coverslip-adhered mLD caps for which TAG droplets were initially spread across the coverslip by a vigorous N_2_ stream in the absence of an aqueous buffer. 3 µL of TAG was pipetted onto the center of a lightly cleaned glass-bottom petri dish prior to a focused N_2_ stream that lightly aerosolized the TAG and dispersed it across the coverslip. Visual inspection of the coverslip was able to readily determine if the small droplets were made across the coverslip (Fig. S2), after which an aqueous buffer was added to the dish (Fig. 3).

**Figure 3:**
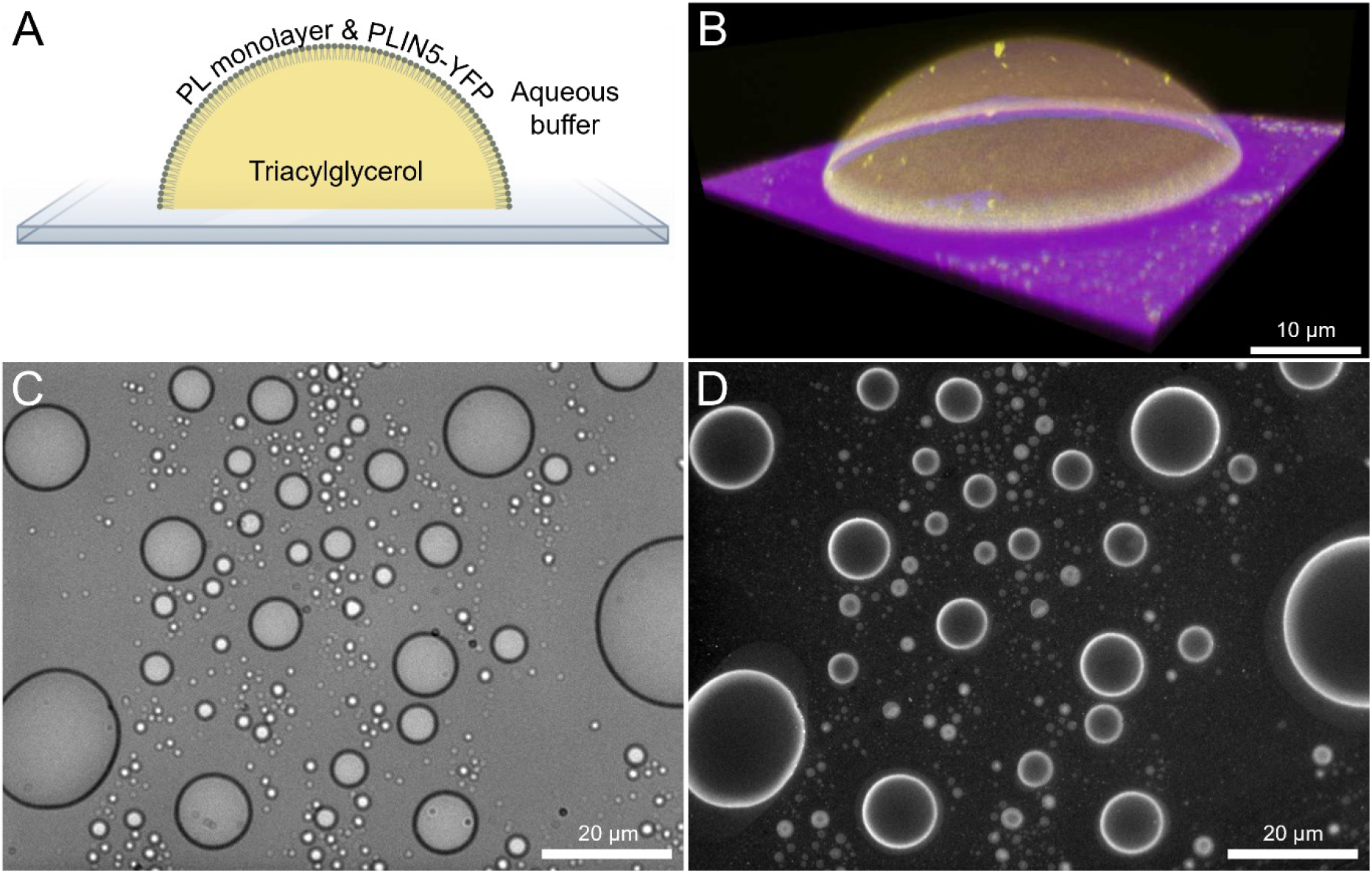
Model lipid droplet caps. (A) Coverslip-bound mLD caps provide optical accessibility by inverted microscopy and exchange of the surrounding aqueous buffer without enclosure with an imaging chamber. (B) Confocal imaging and 3D reconstructions show the contact angle of the droplet to the coverslip as labeled by PLIN5-YFP (*yellow*) and DOPE-Cy5 (*magenta*). Further examples of the near-hemispherical mLD caps are shown in the Supplemental Figures (Fig. S3). (C and D) Many mLD caps may be observed simultaneously with (C) brightfield imaging of the oil-water index contrast or (D) epifluorescence imaging of DOPE-Cy5 on the mLD membrane.

In the absence of surfactants (*i.e*., PL or proteins), a gentle buffer exchange resulted in negligible change to the oil droplet shape for droplets smaller than 80-µm diameter. When PL was present in the aqueous buffer or dissolved within the TAG, only a gentle flow within the aqueous buffer was permitted so as to not perturb the cap shape. In the presence of surfactants and the absence of flow, droplets larger than 40-µm diameter lifted off the glass within 2 hr. Droplets smaller than 40-μm diameter were stably adhered to the coverslip and readily analyzed by long-duration optical or scanning probe methods.

Buffer-covered mLD caps could be readily imaged with confocal (Fig. 3B), brightfield (Fig. 3C), and epifluorescence imaging (Fig. 3D). The coverslip adherence enabled their viewing with inverted microscopes and without concern of a limited objective working distance. Accordingly, the mLD caps are an excellent model system to be used with other fluorescence and scanning probe techniques, as described below.

### Size distributions of mLDs

mLDs were made from <1-µm to >20-µm diameter by altering the TAG:PL ratio (Fig. 4). The TAG:PL ratio determines the volume-to-surface area ratio during sonication prior to sample creaming. Oil-in-water mLDs created via sonication or vortexation and imaged within glass chambers allowed the mLDs to float to the glass slide and expose their underside for unobstructed viewing. Floating mLDs of varying sizes could be observed at differing imaging focal planes at the time of image acquisition (Fig. S1). The smallest droplets were found at a higher focal plane and closer to the glass slide than the larger droplets. To assess the sizes of the mLDs in each sample, a focal plane that corresponded to the widest part of majority of the droplets was used (Figs. 2B, C and 4).

**Figure 4.**
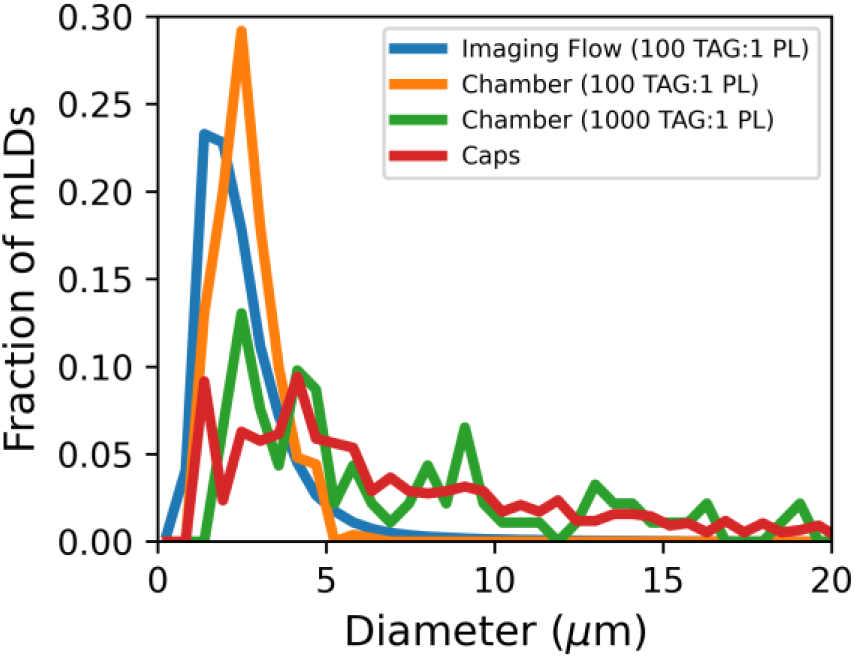
Size distribution of model lipid droplets. Varying the TAG:PL ratio and the imaging method determines the size of the mLDs observed. Imaging flow cytometry enables observing >100k buoyant mLDs within 3 min. Trapping buoyant mLDs between a coverslip and glass slide chamber enabled inverted microscopy imaging. The mLD caps provided the widest distribution of mLD sizes for simultaneous observation.

The air objectives of imaging flow cytometry provided a low numerical aperture and a deep depth of field. Image culling based on the image contrast was used select only those in focus. The droplets that triggered the cytometer were imaged, and the final data set was created by culling the images based on the droplet size, shape, and number, as described in the Methods. The lower NA of the imaging flow cytometer objective yielded lower image resolution and difficulty in distinguishing if more than one mLD had aggregated together. This complexity has limited our ability to study mLDs with proteins via imaging flow cytometry because the proteins caused the mLDs to flocculate.

The mLD caps provided the widest range of droplet sizes for analysis. Varying the coverslip treatment could alter the adhesion of the caps to the glass and the contact angle of the droplet. Typically, surface treatments were used that provided hemispherical caps (Fig. S3). Both the buoyant droplets in glass chambers and the mLD caps provided the ability to perform long-duration imaging on individual droplets.

### PL binding and exchange

The formation of the PL monolayer on the mLD caps was monitored by the increase in PL monolayer fluorescence versus time (Fig. 5A). LUVs were created with varying non-fluorescent PL compositions and either 1 mol% Cy5 or 1 mol% TF. The LUVs were added to the oil-on-glass caps at 0.4 mg/mL in IB and epifluorescence images were taken every 5 min. To observe the monolayer brightness distinct from the unbound LUVs in solution, the unbound LUVs were rinsed away, images were taken, and LUVs were added back. The first 20 min of LUV exposure occurred with Cy5-labeled LUVs and monolayer brightness was apparently saturated within 5 min. To test the LUV exchange or addition of more PL to the monolayer after 20 min, the Cy5 LUVs were removed, and TF LUVs were added. The non-fluorescent PLs were 100 mol% PC; 73 mol% PC and 27 mol% PE; or 65 mol% PC, 27 mol% PE, and 8 mol% PI. Both the binding of Cy5 LUVs to the bare oil-water interface and the binding of TF LUVs to Cy5-coated mLD were enhanced when the non-fluorescent lipids included PE and PI rather than only PC.

**Figure 5.**
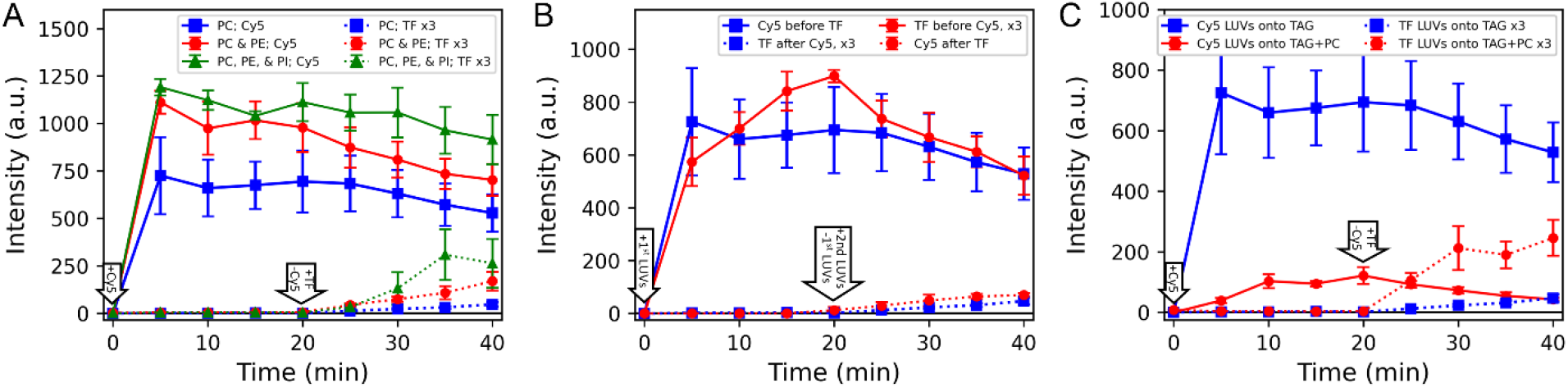
LUV binding to mLDs. Coverslip-adhered mLD caps enable time-dependent measurements of PL binding to the mLD monolayer. (A) The binding of LUVs was measured through the sequential addition of Cy5-labeled LUVs followed by TF-labeled LUVs. The Cy5-labeled LUVs seemingly saturated the monolayer within 5 min when present at 0.4 mg/mL surrounding the caps. After 20 min, the Cy5 LUVs were replaced with TF LUVs to monitor the continued binding or exchange of PL to the caps. The TF signals shown here are magnified by 3x to better visualize the dimmer emission from TF compared to Cy5. (B) The results are similar if the Cy5 or TF-labeled LUVs bound to the mLD caps first. The nonfluorescent PL was PC in the LUVs. (C) The PL monolayer created by the inclusion of 4% w/w PC dissolved within the TAG resulted in inhibition of PL binding or exchange from LUVs as compared to when the TAG had no PL dissolved within it.

The control experiment was performed in which the mLD caps were exposed to TF-labeled LUVs prior to the Cy5-labeled LUVs with indistinguishable results regarding the reduced but notable binding of the second type of LUVs (Fig. 5B). A potentially important distinction is that the fluorescence intensity of the TF-labeled LUVs did not saturate within 20 min as did the Cy5-labeled LUVs. This result is more consistent with the long durations required for LUV binding for a saturated reduction in the monolayer surface tension, as described below. In all cases, for the consistent imaging parameters used here, the TF fluorescence emission was approximately one third as intense as the Cy5 fluorescence emission, thus the TF signals plotted are multiplied by three for clarity on the same axis.

The binding of LUVs to mLD caps that included PC dissolved within the TAG appeared consistent with the binding of LUVs to mLDs that were previously exposed to LUVs (Fig. 5C). In all cases, when the Cy5 LUVs were exchanged for TF LUVs, the mLD monolayers decreased in Cy5 brightness suggesting an exchange of PL between the mLD monolayer and the LUVs. Control experiments confirmed that the decrease in Cy5 brightness was not due to photobleaching of the Cy5, which is expected to be less than 7% over the course of these experiments. This result suggests that the binding and exchange of PL between the mLD monolayer and LUVs in solution is independent of whether the monolayer was formed by prior LUV binding or dissolution of PL form the mLD TAG interior. Cy5 PLs on the monolayer, and perhaps all the PLs on the monolayer, were able to exchange with the LUVs in solution. The mLD monolayers seemingly approached the composition of the surrounding LUVs regardless of the initial monolayer composition.

### PLIN5 diffusion on mLDs

The diffusion of PLIN5-YFP on mLD caps and buoyant mLDs were observed with FRAP and FCS, respectively (Fig. 6). PLIN5-YFP was bound to mLD caps in which the PL monolayer was created by 20-min of LUV binding, PLs initially mixed within the TAG, or with no PL. The FRAP recovery curves yielded the PLIN5-YFP diffusion rate and the mobile fraction of PLIN5-YFP. The PLIN5-YFP recovery time constants were 14 ± 5, 16 ± 4, and 22 ± 11 sec, respectively, yielding diffusion coefficients of 0.21 ± 0.13, 0.20 ± 0.05, and 0.14 ± 0.10 μm^2^/s, respectively, for the three PL conditions; uncertainties are reported as the standard deviations of the repeated measurements.

**Figure 6.**
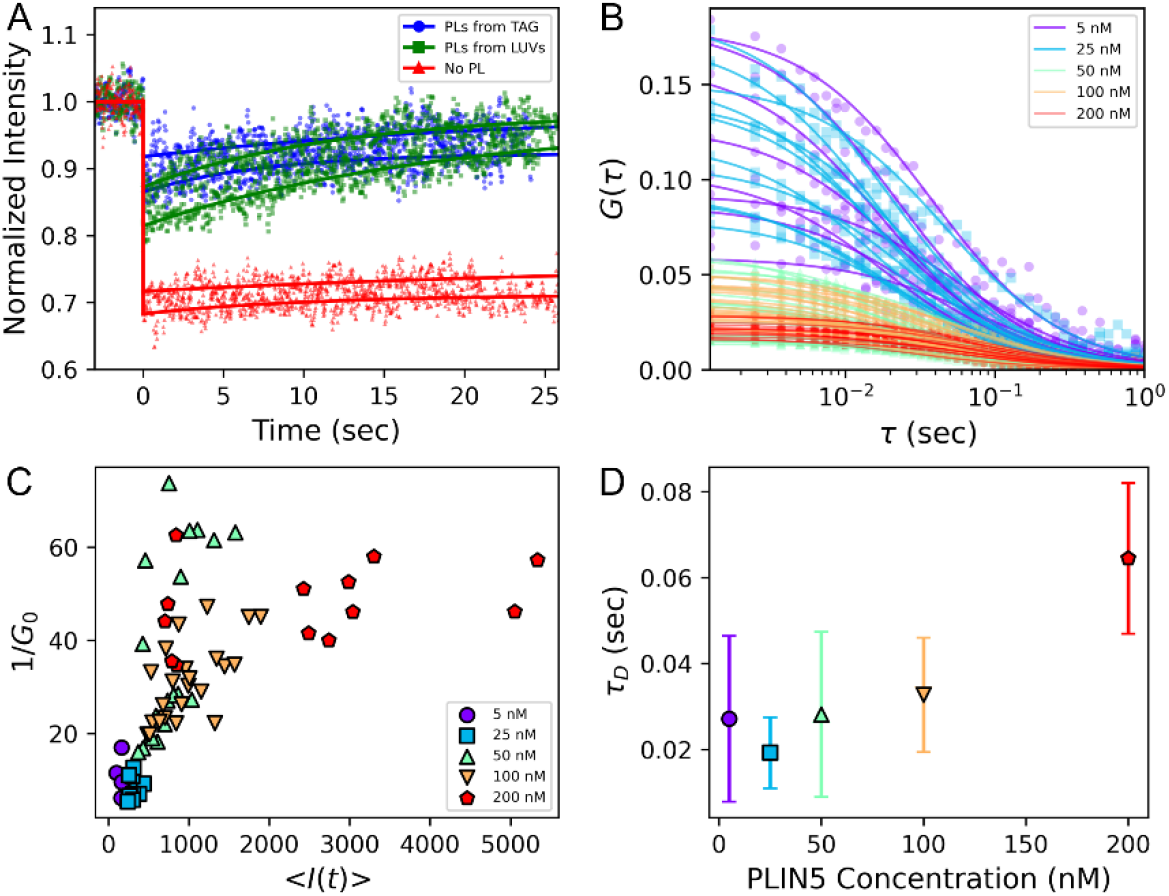
Diffusion of PLIN5-YFP on mLDs. The diffusion of PLIN5-YFP on the mLD membranes depends on the PL present and the PLIN5-YFP concentration. (A) The coverslip-adhered mLD caps allow for the diffusion measurements of mLD-bound PLIN5-YFP via FRAP with two repeats shown per condition shown. The 4 sec bleaching duration is not shown. The diffusion of PLIN5-YFP varied with how the PL monolayer was created. Most notably, in the absence of PLs, the PLIN5-YFP diffused slower and was seemingly immobile on the mLD. Experimental data points (*symbols*), fits to Eq. 1 (*lines*), and representative images (Fig. S4) are shown. (B) Autocorrelations of PLIN5-YFP at varying solution concentrations (*symbols*) were well fit by the expected 2D Browning diffusion function (*lines*, Eq. 3). (C) The inverse of *G_0_* and the intensity of detected fluorescence emission both correspond to the concentration of diffusers present within the observation volume. For intermediate concentrations of PLIN5-YFP, the inverse of *G_0_* and <*I*(*t*)> are well correlated. (D) The observed dwell time of PLIN5-YFP was consistent for all PLIN5-YFP concentrations ≤100 nM with a median value of 27 ± 16 ms. When [PLIN5-YFP] = 200 nM, the observed dwell time was 64 ± 18 ms. Uncertainties are the standard deviations of measured values.

The mobile fraction of PLIN5-YFP varied with the mLD monolayers. In the absence of PL, the PLIN5-YFP bound to the mLD demonstrated little free diffusion with the FRAP curve negligibly recovering (Figs. 6A, S4). Without PL, the bleached spot maintained prominence with (12 ± 3)% of the fluorescence recovering. With PL, (60 ± 18)% and (86 ± 1)% of the fluorescence recovered when the PL came from the TAG or LUVs, respectively. Accounting for diffusion during the 4 sec of bleaching, the maximum recovery fraction of PLIN5-YFP was (94 ± 3)%, (97 ± 1)%, and (62 ± 5)% These results suggest that PLIN5-YFP mobility is highly dependent on the PL monolayer coating the mLDs.

For FCS, PLIN5-YFP was added to the spherical, buoyant mLDs prior to their incorporation into the microsphere-supported glass microscopy chambers. The diffraction-limited point of observation on the mLDs was the bottom of the spherical mLDs that was closest to the inverted microscopy objective. By focusing on the bottom point of the mLD, the focused laser excitation and the emission transmitted to and from the microscope objective without refraction through the curved mLD surface. Additionally, the bottom of the floating mLDs was >5 μm from the potentially perturbing coverslip. Increasing the solution concentration of PLIN5-YFP resulted in a higher concentration of LD-bound PLIN5-YFP, as assessed by both the increased brightness of the PLIN5-YFP bound to the mLD surface and the reduced amplitude of the PLIN5-YFP autocorrelation (Fig. 6C). The linear correlation between 1/*G_0_* and the emission intensity observed was readily apparent for intermediate emission intensities, which was only observed for [PLIN5-YFP] ≤ 100 nM. When [PLIN5-YFP] exceeded 100 nM in the surrounding solution, the brightness of the PLIN5-YFP on the mLD membrane continued to increase, but the amplitude of the autocorrelation did not continue decreasing. This is consistent with the highest PLIN5-YFP concentrations demonstrating a changed binding modality and slowing of the PLIN5-YFP diffusion, such as homo-oligomerization or aggregation. The characteristic dwell time of the PLIN5-YFP remained relatively constant for [PLIN5-YFP] ≤ 100 nM and *D* = 0.23 ± 0.02 μm^2^/s. But PLIN5-YFP demonstrated slower diffusion at 200 nM with *D* = 0.01 ± 0.003 μm^2^/s (Fig. 6D).

### Phospholipid monolayer packing measured by FLIM

The fluorescence lifetime of Flipper-TR reports the packing of the PLs within a membrane^27, 28^. Shorter lifetimes reflect a lower density or more fluid PLs as the Flipper-TR has more molecular flexibility for quicker relaxation. FLIM images of mLD caps coated with non-fluorescent PLs and Flipper-TR show that the Flipper-TR is concentrated in the monolayer surrounding the mLD and has minimal partitioning into the bulk oil or bulk aqueous phase (Fig. 7A).

**Figure 7.**
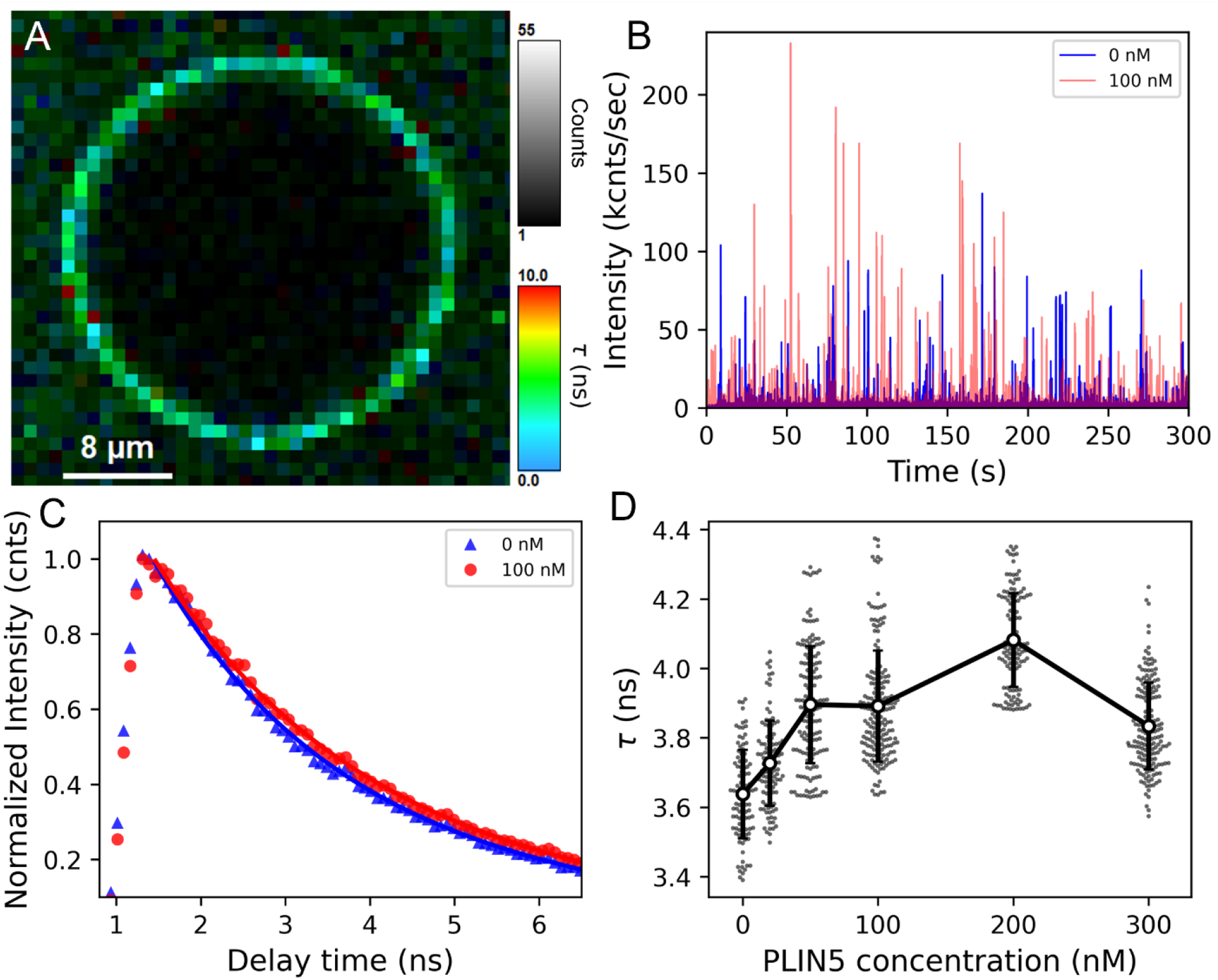
Fluorescence lifetime imaging microscopy on mLD membranes. The fluorescence lifetimes of Flipper-TR were observed within the monolayer of mLDs. (A) FLIM imaging of coverslip-adhered mLD caps showed that Flipper-TR concentrates in the PL monolayer surrounding the mLDs. (B) Representative intensity versus time data showed the bursts of emission as nanoscale mLDs diffused through the detection volume in the presence of 0 or 100 nM PLIN5. (C) Representative photon delay time histograms (*symbols*) from Flipper-TR on mLDs in the presence of 0 or 100 nM PLIN5 that were fit to a biexponential decay (*lines*). Note the longer decay time for 100 nM versus 0 nM of PLIN5. (D) The increase in emission decay time upon the addition of PLIN5 suggests that PLIN5 incorporation causes the PLs to become more tightly packed. The swarm plot of individual data points (*gray*) represents the distribution of decay times with their mean and standard deviation overlayed (*black*).

The monolayer packing was measured both by imaging mLD caps and by observing the transient bursts of small mLDs diffusing through a fixed observation spot. The mLD caps provide the changes versus time for a single mLD, but they are vulnerable to photobleaching, and the slow image acquisition rates by FLIM impose challenges to observe many mLD caps. Alternatively, by observing the spontaneous diffusion of small mLDs through a fixed observation spot, many mLDs were observed within a single 3-min acquisition, and photobleaching was less of a concern since a single mLD is unlikely to be observed twice. The buoyant nanoscale mLDs were sufficiently stable when initially dispersed throughout the 20 μL sample for observation over 20 min with multiple 3-min acquisition times. This is supported by the expected balance of the droplets’ buoyant force and drag coefficient yielding an expected terminal velocity of 4 nm/s for 0.3-μm diameter droplets and 200 nm/s for 2-μm diameter droplets within the 1.4-mm tall hemispherical sample of IB. The thermal diffusion rate of the 0.3 and 2-μm diameter droplets is expected to be 1.4 and 0.2 μm^2^/s, respectively, yielding a transit time of 0.12 and 0.8 s, respectively, through the diffraction-limited observation volume, which is far greater <10 ns fluorescence decay time. Accordingly, the dynamic small mLDs suspended within the IB are well suited for FLIM observation despite their buoyancy and thermal diffusion.

The fluorescence lifetime of Flipper-TR was observed in small mLDs upon the addition of non-fluorescent PLIN5 (Fig. 7). PLIN5 bound to the mLDs, intercalated between the PLs, caused increased PL packing, reduced Flipper-TR molecular flexibility, and increased the Flipper-TR fluorescence lifetime. The acquired data shows modest changes in *τ* with variability depending on the data fitting methods employed.

Sample-to-sample reproducibility demonstrated a maximum of 1.1% variation in the mean *τ* observed upon application of the selection criteria described in the Experimental Procedures. When the data from all samples, including those with low total counts or unusually large burst of emission, were incorporated, a maximum of 9.6% sample-to-sample variation was observed. The variable fitting methods employed demonstrated a standard deviation of fits less than 3.1% of the mean for all conditions. A 7.5% increase in *τ* for [PLIN5] ≥ 100 nM versus [PLIN5] = 0 nM and a propagated uncertainty of 3.3% yields a Z-score of 2.3 and a P-value of 0.01; *τ* significantly increased with the addition of ≥100nM PLIN5.

### Droplet surface tension changes

The surface tension on the mLDs determines the energetic cost of increasing the droplet area. Force spectroscopy with an AFM was used to measure the force necessary to deform mLD caps and increase their surface area. We modeled the mLD cap shape initially as a hemisphere and assumed a constant droplet-to-coverslip contact area, a constant droplet volume, and a minimum droplet area as the AFM tip pressed upon the cap (Eq. 5). Numerically solving for the droplet shape given the above constraints as the cap was compressed by the AFM yielded the estimated surface tension with droplet height compression (Fig. 8B, C). For a few percent change in the droplet height, the force upon the AFM tip was estimated to be proportional to the height change of the droplet with a slope of -(4.33 ± 0.01)γ (Eq. 6, Fig. 8D).

**Figure 8.**
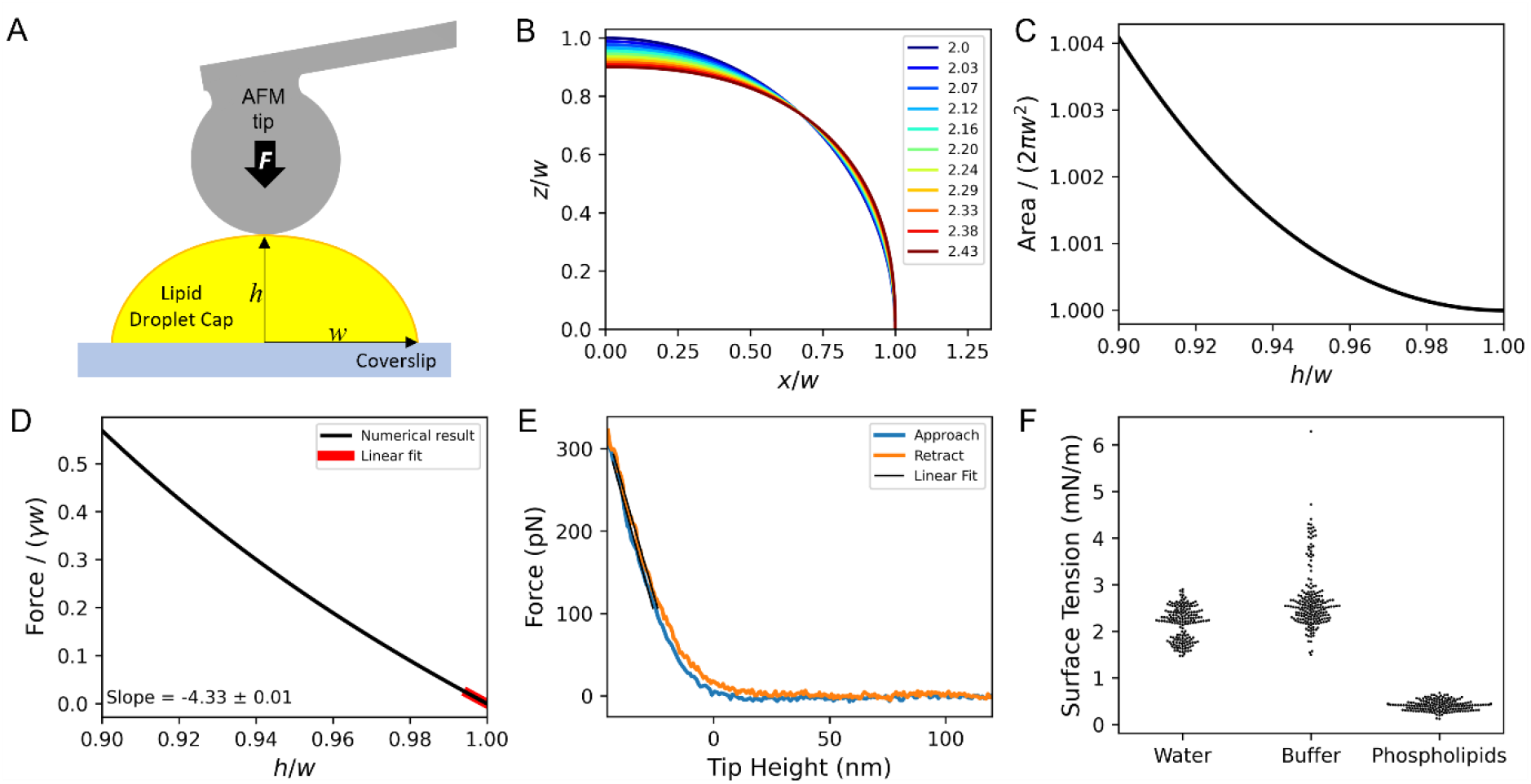
Measuring membrane tension with AFM. (A) AFM force spectroscopy of mLD caps with a colloidal tip measures the force of increasing the mLD cap area. We assumed the mLD maintains a constant volume and constant interaction area with the glass coverslip as it is compressed to a shorter height. (B) The mLD cap shape was approximated as smoothly transitioning from a hemisphere towards a cylinder (Eq. 5) with a constant width (*w*), constant volume, changing height (*h*), and changing surface area. The exponential value *n*, as shown by the line color, was numerically determined to fit these constraints (Fig. S5). (C) The fractional increase in the cap area is determined by the cap’s height-to-width ratio. (D) The force on the AFM tip is proportional to the change in the mLD cap area, which is approximately linear for the first 100 nm compression of a 20-µm-wide cap. (Eq. 6). (E) The linear force-to-indentation depth relationship was experimentally observed upon approach and retraction of the AFM tip on mLD caps. (F) Repeated measurement on mLD caps composed of 97% TAG demonstrated that the cap surface tension was indistinguishable when the cap was surrounded by water or IB, but the presence of PLs resulted in an order-of-magnitude decrease in the cap surface tension. These mLD caps were formed and stabilized for >5 hr prior to measurement.

The experimental approach and retraction curves showed a consistent force upon the AFM tip versus tip height (Fig. 8E). At far separation distances, there was no measured force upon the AFM tip, which shows the minimal drag on the AFM tip as it was moved through the buffer. The first forces upon the AFM tip were presumably non-contact forces independent of the mLD surface tension. After the cap was compressed >20 nm, a linear force versus height was observed resisting compression of the cap, which agreed with our numerical model. The slope of the force versus height was determined and fit to Eq. 6 to calculate the surface tension of the mLD cap. Repeated measurements on individual caps, between caps in a single dish, and in separate dishes yielded consistent surface tensions for the three conditions tested while using the 97% TAG. The surface tension decreased from 2.2 ± 0.4 mN/m and 2.5 ± 0.6 mN/m in Milli-Q water and IB, respectively, to 0.4 ± 0.1 mN/m in 0.8 mg/mL LUVs after equilibrating for 1 hr (Fig. 8F); the average values are reported as the median and standard deviation of all force curves for each condition.

Our custom pendant droplet tensiometer employed a cuvette and needle to minimize both the droplet volume and the surrounding buffer volume. The 30-gauge blunt needle provided an internal diameter of 160 μm that was suitable to hold droplets up to 10 μL within the disposable polystyrene microcuvette. Under conditions of <10 mN/m surface tension, <6 μL droplets were necessary to limit detachment. Accurate tension measurements could be made on droplets >1 μL. Inserting the 90° curved needle into the bottom of the microcuvette such that the tip of the needle was within 2 mm of the bottom of the cuvette allowed for the droplet to be fully submersed within 300 μL of buffer; we typically used 400 μL of buffer to ensure the droplet was >2 mm from the buffer-air interface. The 4.5 mm wide cuvette provided 0.9 mm of space between the sides of the cuvette and the 10 μL droplets.

Our custom optical setup incorporates standard optical components with a 4x objective to provide brightfield images of the droplet shape (Fig. 9A). The software OpenDrop automatically detects the droplet’s edge and fits it to the Young-Laplace equation to report the droplet’s volume, surface area, and tension for each frame of the timelapse movie. A computer-controlled syringe pump was used to grow the oil droplet within the cuvette containing the aqueous buffer. The tension could not be accurately determined until the droplet grew sufficiently large to reveal a non-spherical shape.

**Figure 9.**
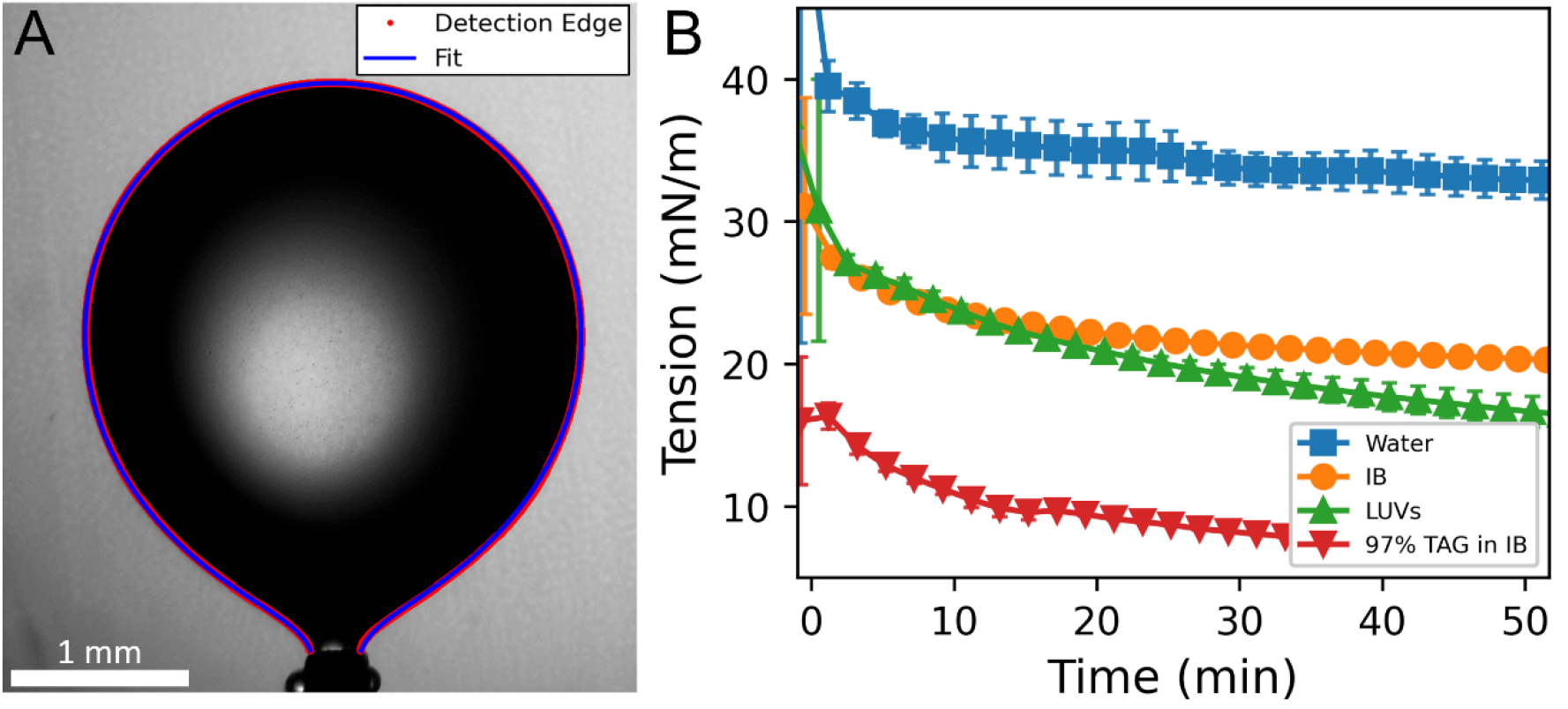
Pendant droplet tensiometry of model lipid droplets. A custom pendant droplet tensiometer has enabled the measurement of the TAG-buffer interfacial tension versus time upon the addition of buffers or PLs. (A) Droplet images were analyzed with OpenDrop identify the droplet edge and fit it to the Young- Laplace equation. (B) The tension versus time shows the sorting of surfactants from the aqueous buffer or from the TAG to decrease the droplet tension after its formation (Fig. S6). When >99% pure triolein was used, the triolein-in-water tension well matches prior reports. The addition of salts in IB or 0.8 mg/mL PC LUVs decreased the tension. Impurities within 97% triolein demonstrated a significantly lower tension than the other conditions.

The surface tension of the droplets decreased versus time as the surfactants within the oil or buffer concentrated on the oil-water interface. The 99% triolein in Milli-Q water demonstrated a surface tension of 33 ± 1 mN/m while surfactants within the IB resulted in an observed surface tension of 20 ± 1 mN/m after 50 min of equilibration. The surfactants present as impurities within the 97% triolein resulted in a rapid decrease in its surface tension in IB to 7.8 ± 0.4 mN/m over the 40 min of observation. The presence of PL as LUVs in the surrounding buffer resulted in a consistently decreasing surface tension that was expected to decrease below the observed 16.5 ± 1 mN/m should there have been greater time for the LUVs to bind and saturate the monolayer.

## Discussion

Testing the biophysics of lipid droplets requires novel methods capable of isolating and measuring the dynamic properties of the oil-water interface. This interfacial membrane is challenging to measure optically because it typically consists of a curved boundary with a high index of refraction contrast that distorts imaging through the droplet and complicates analysis of on-membrane diffusion. Further, the low mass density of the LD oily interior causes the droplets to be buoyant in aqueous buffers and potentially float out of the working distance of inverted microscopy objectives. Through innovations in sample confinement, surface adhesion, and imaging flow cytometry, we have enabled high-resolution imaging of LDs with exchangeable aqueous buffers.

Transient adhesion of TAG droplets to microscopy coverslips was achieved to yield a moderately lipophilic surface for the creation of mLD caps. mLD caps provide the experimental benefits of allowing for aqueous exchange surrounding the mLDs with a planar bottom while maintaining proximity of the mLDs to the short working distance, oil-immersion inverted microscopy objectives. These caps were used to measure the binding rate and the diffusion of PL and proteins on the mLD surface. Additionally, mechanical deformation of the caps with an AFM scanning tip has enabled measurement of the mLD surface tension.

The creation of mLDs through sonication, vortexing, or N_2_ fragmentation creates heterogeneous, micron-sized droplets. This size distribution is both a feature and a challenge for systematic mLD analysis. Nanoscopic mLDs with a low TAG:PL ratio were measured with FLIM when initially mixed throughout the aqueous buffer and prior to their floating out of view. Larger mLDs were confined within microbead-stabilized glass chambers to keep the mLDs within the short working distance of the oil-immersion objective. These chambers enabled long-duration, single-spot observation of membrane-bound molecules diffusing on the membrane by focusing on the bottom of a spherical mLD. Finally, the surface-adhered mLD caps were measured during exchange of the aqueous buffer for time-dependent measurements of the membrane change.

The monolayer surface tension is among the most important physical properties for governing the activity of LD-associated proteins for LD nucleation, growth, and the regulation of lipolysis. We employed three methods of measuring the LD surface tension: AFM force spectroscopy, FLIM, and pendant droplet tensiometry. These three methods provide distinct physical methods to measure the PL monolayer tension. The scanning AFM probe compressed the mLD cap and measured the force resisting expansion of the cap area that is directly related to its surface (Eq. 6). Successful AFM force spectroscopy required a passive probe-membrane interaction without significant binding, adhesion, or hysteresis. Should the tips crash into the droplet and become coated with TAG, a rigorous rinsing of the used tips in isopropyl alcohol proved to restore the tips to their original, clean state for negligible interaction with the droplet upon repeated use. Commercial cleaners such as Hellmanex III to clean the AFM probe resulted in a detachment of the colloidal microsphere from the AFM cantilever.

Commercial pendant droplet tensiometers are readily available, however, we aimed to create a cost effective, precise, custom tensiometer for samples of minimum volume. Our system incorporates a semi-micro cuvette with a 30-gauge needle inserted near its bottom. The sample enables viewing of 1 – 10 µL oil droplets within 300 µL of aqueous buffer. This small volume of aqueous buffer will be especially important as we use the system to reduce the consumption of expensive reagents on the mLD surface in future studies. The optical setup necessary to resolve the shape of 2-mm diameter droplets may be as simple as a consumer camera with a macro lens.^29, 30^ Further, the use of open-source software for image analysis (*i.e.*, FIJI and OpenDrop) allows for custom pendant droplet tensiometers to be a cost-effect experimental setup.^31, 32^

FLIM offers a technique to optically measure membrane packing without physically manipulating nor precisely resolving the LD shape. The fluorescence lifetime of membrane-embedded fluorophores is difficult to correlate to physical measurements of membrane tension; however, the fluorescence lifetimes of Flipper-TR is correlated with PL packing, composition, temperature, phase, and tension.^27, 33^ Our discovery that Flipper-TR may be concentrated in the LD monolayer compared to the bulk oil or aqueous phases in the samples enables the use of FLIM to resolve membrane changes coupled with metabolic processes. The demonstration that the Flipper-TR fluorescence lifetime increases upon the addition of PLIN5 reveals that PLIN5 binding is affecting the packing of the PL on the LD membrane via mechanisms and with ramifications that are yet to be discovered.

The LD surface tension depends on the surfactants present. Natural surfactants, such as PLs, segregate to the oil-water interface, but kinetic limitations in LUV binding complicate the creation of the PL monolayer from LUV fusion (*i.e*., Fig. 9B). Dissolving the PL within the bulk TAG prior to mLD or pendant droplet formation is thought to create a complete monolayer quickly,^21^ but a detailed analysis is warranted in future studies. Interestingly, the diffusion of mLD-bound PLIN5-YFP was weakly dependent on the source of the PL, but PLIN5-YFP diffusion was inhibited if no PL was present (Fig. 5B). We hypothesize that the incomplete PL monolayer created by <1 hr of LUV binding is sufficient for natural PLIN5-YFP diffusion, but future studies are warranted to establish if PL density on the mLD affects other properties of protein behavior, such as the maximum bound protein density or protein-protein interactions. Comparing PL monolayers created by LUV binding versus from PL dissolved within the TAG will further elucidate how protein behavior is affected by the PLs.

The surface-active molecules for mLDs may come from a variety of sources, including from the buffer and from impurities within the TAG used (Fig. 9). IB was created by mixing salts with Milli-Q water, and included surfactants that reduced the surface tension of otherwise pure TAG droplets in the pendant droplet tensiometer. The 97% triolein and 99% triolein used here demonstrated vast differences in the surface-active impurities present. The surface-active impurities from the 97% triolein prevented AFM force spectroscopy from identifying the differences between Milli-Q water and IB (Fig. 8). We expect that dissolving PL within the TAG prior to droplet formation will result in a quickly saturated monolayer of PL with a tension <1 mN/m and similarly mask any differences between Milli-Q water and IB. During N_2_-stream fragmentation and buffer exchange of mLD caps, we have qualitatively observed the dramatically reduced surface tension of the oil-water interface when PL is previously dissolved within the TAG. The methods developed here will be the basis of our future exploration of PL-dependent protein behaviors.

This manuscript reports the development of methods for creating and analyzing the biophysical properties of mLDs. We have identified the challenge of observing curved, buoyant mLDs with conventional inverted microscopy methods. Through the confining of micron-sized mLDs within microbead-supported glass chambers, the bottom of the buoyant mLDs could be observed without refractive distortion by the LD surface or the disruptive proximity of a substrate. Sub-micron droplets were measured by FLIM with a stationary observation volume while they diffused within the aqueous buffer. To study the effects of aqueous buffer exchanges and to test the rapid dynamics of protein or ligand binding kinetics, substrate-adhered mLD caps demonstrated long-term stability during buffer exchange while maintaining convenient high-resolution imaging by oil-immersion objectives. While complemented by scanning probe manipulation, these mLD caps have the potential be a near-universally applicable model system for the studying of the monolayer biophysical properties. The collection of cross-validating methods presented overcame numerous prior experimental challenges and have been used to measure the binding, diffusion, sorting, and tension changes of mLD monolayers.

## Experimental Procedures

### Resource Availability

Further information and requests for resources and reagents should be directed to and will be fulfilled by the Lead Contact, Christopher V. Kelly. The custom analysis code generated during this study are available on GitHub.com.^34^

### Phospholipid vesicle preparations

Non-fluorescent and fluorescent phospholipids (PLs) were obtained from Avanti Polar Lipids and used without further purification. The nonfluorescent PLs included 1,2- dioleoyl-sn-glycero-3-phosphocholine (PC); 1,2-dioleoyl-sn-glycero-3- phosphoethanolamine (PE); and L-α-phosphatidylinositol (PI). An LD-mimicking PL monolayer membrane was composed of PC (65 mol%), PE (27 mol%), and PI (8 mol%). Fluorescent lipids include 1,2-dioleoyl-sn-glycero-3-phosphoethanolamine-N-Cyanine 5 (Cy5), 2-distearyl-sn-glycero-3-phosphoethanolamine-N-TopFluor AF488 (TF), and Flipper-TR (Cytoskeleton, Inc.).

To make multilamellar vesicles (MLVs), the PLs were mixed in chloroform, dried under a nitrogen stream, and kept under vacuum for >30 min. The dried PL film was hydrated in intracellular buffer (IB) and vortexed to yield MLVs. IB consists of 10 mM HEPES, 140 mM KCl, 6 mM NaCl, 1mM MgCl_2_, and 2 mM EGTA in Milli-Q water at pH 7.4. Large unilamellar vesicles (LUVs) were created by extruding MLVs through 100-nm diameter pores 21 times. Fluorescent Cy5 or TF, if used, were mixed in chloroform with the non-fluorescent PLs prior to making MLVs. When used, Flipper-TR was added to LUVs by mixing 500 μL of 0.1 g/L non-fluorescent LUVs in IB with 0.6 μL of 1 mM Flipper-TR in DMSO and immediately vortexing to facilitate the incorporation of the Flipper-TR into the LUVs prior to its aggregation.

### Droplet-embedded vesicles (DEVs)

As described previously, GUVs were made through electroformation^35^ and DEVs were made by mixing TAG droplets with the GUVs^22^. To form the GUV growth chamber, 180 μg of the LD-mimicking PL mixture with 1 mol% Cy5 for >30 min was dried under vacuum on ITO plates. The growth chamber was assembled with two ITO plates separated by a silicon spacer, filled with 0.75 mL of 200 mM sucrose, and subjected to a 10 Hz, 4 V_pp_ sine wave function generator for 3 hrs at 50°C. For osmotic balance, TAG droplets and GUVs were diluted in a 66% IB and 33% Milli-Q water mixture. TAG droplets were created by mixing 5 μL of >99% pure glyceryl trioleate (T-235, NuChek Prep) with 70 μL of diluted IB, vortexing for 60 sec, and bath sonicating for 10 sec until a cloudy mixture was observed. The TAG droplets were kept on a rotator to minimize coalesce and used within 2 hrs. DEVs were created by mixing 10 μL of GUVs with 20 μL of the TAG droplets and storing them on the rotator for 30 min at RT. The DEVs were imaged in a glass-bottom 394-well plate on the Dragonfly confocal microscope by mixing 5 μL of the DEVs with 40 μL of diluted IB and either 0 or 190 nM PLIN5-YFP.

### Buoyant model lipid droplets (mLDs)

Buoyant, spherical mLDs were typically formed by mixing MLVs with glyceryl trioleate as the model TAG. Typically, the buoyant mLDs were created through bath sonication for 30 min (Ultrasonic Cleaner 96043-936, VWR Symphony). Varying the TAG-to-PL ratio altered the final mLD size; however, excessive sonication also produced mLDs with incomplete PL covering and emulsion creaming. The mLDs were stored on a laboratory rotator at 4°C, which was essential for keeping the buoyant mLDs well mixed through the aqueous buffer. mLDs >1-μm diameter were used or discarded the same day they were generated. mLDs imaged within microscopy chambers were mixed with 20-µm diameter nonfluorescent polystyrene bead spacers (74491, Millipore Sigma) were added prior to the sample confinement between a glass slide and a coverslip. The glass slides and coverslips were previously rinsed with water, ethanol, dried under N_2_ stream, and passivated with a casein. Passivation included exposure to 20 mg/mL of casein (218682, Sigma-Aldrich) for >20 min prior to thorough rinsing with Milli-Q water.^25^ The buoyant mLDs floated to the glass slide, while the coverslip was closer to the inverted microscope objective (Fig. 2A). The bottom of the mLDs was the focus for our FCS observation point such that neither the illumination nor emission light refract through the mLDs on its path to or from the objective.

The buoyant mLDs were labeled with proteins or Flipper-TR and stored on the rotator prior to their confinement within the imaging chambers. Flipper-TR was incorporated by mixing the mLDs with Flipper-TR-labeled LUVs, as described above.

### mLD Caps

Glass-supported mLD caps were created by using a regulated adhesion between the TAG and the glass coverslip. On a glass-bottom 35 mm dish (MatTek) or a glass-bottom 96-well plate (Cellvis), the glass surface was cleaned with water, surfactants, ethanol, and dried with a N_2_ stream. The glass-bottom 96-well plates required 2% Micro90 cleaning for >10 min to remove residue from the manufacturer. The 35 mm dishes required no surfactant cleaning. Excessive surfactant cleaning or any plasma cleaning (Harrick Plasma Inc.) resulted in insufficient oil-glass interaction and detachment of the droplet caps from the coverslips. A more hydrophobic glass treatment (*i.e*., coverslips without surfactant cleaning) yielded more stable and flatter caps. Glyceryl trioleate was either >99% purity (T-235, NuChek Prep) or 97% purity (CAS: 122-32-7, Sigma Aldrich). When dissolving PL within the TAG prior to creating the caps, PL in chloroform was added to the TAG, and the chloroform was evaporated under vacuum for >4 hr for a final PL concentration was 2% w/w. The TAG was added by pipetting 3 μL of triolein on to the center of the coverslip prior to a vigorous N_2_ stream that caused a fragmentation of the oil into small droplets randomly scattered across the coverslip (Fig. S2). Visual inspection identified the small oil droplets adhered to the coverslip when sufficient spreading of the oil was achieved, and aqueous buffers could be gently applied to coat the caps. The mLD caps were sufficiently stable to accommodate numerous buffer exchanges, however the caps were more likely to detach from the coverslip and float in the buffer if PLs were present. As shown in the data presented here, the caps were consistently stable during buffer exchanges and time-lapse imaging lasting >2 hr. Non-fluorescent PLIN5 was added after sonication and stored for 90 minutes on the rotor prior to FLIM measurements.

### Protein purification and preparation

Baculovirus for protein expression were prepared using the Bac-to-Bac expression system (Invitrogen). pFastBac1 constructs for PLIN5-yellow fluorescent protein (YFP)-His tag (all mouse proteins) were made using standard molecular biological methods and were confirmed by sequencing. Bacmid DNAs were generated by transformation of DH10Bac E. coli (Invitrogen) with FastBac plasmids following the manufacturer’s protocol. Initial baculovirus stocks were generated by transfection of Sf9 insect cells with bacmid DNA using Cellfectin II reagent (Invitrogen) according to the manufacturer’s protocol, then were amplified by infection of Sf9 cells with the initial baculoviral stocks.

Amplified baculoviral stocks were used to infect High Five insect cells and cells were collected 48 hr after start of infection when cells were 80% viable. High Five cells expressing proteins were lysed by sonication in IB buffer containing 20 ug/mL leupeptin and pepstatin A. For PLIN5-YFP-His tag preparations, 0.5% FOS-CHOLINE-12 detergent (Anatrace) was added to sonicated cell extracts prior to centrifugation at 10,000g. Supernatants were incubated with TALON Cobalt beads (Takara) for 2 hrs at RT and the proteins were eluted from beads using 50 mM sodium phosphate buffer with 150 mM imidazole at pH 7.4. Imidazole was removed from proteins by successive rounds of concentration and redilution with IB buffer using Centricon centrifugal filter units. Protein concentrations were determined by BCA (Pierce), then protein purity was determined from Coomassie-stained PAGE gels.

### Optical Microscopy

Wide-field fluorescence and brightfield images were taken on an inverted microscope (IX71, Olympus) with a 40x, oil-immersion objective and an EMCCD camera (QiClick, QImaging). Confocal fluorescence images were taken with an inverted microscope (DMi8, Leica) with spinning disc apertures (Dragonfly 200, Andor) equipped with a 60x, oil-immersion objective and a sCMOS camera (Zyla, Andor).

### Imaging flow cytometry

Imaging flow cytometry (ImageStream Mark II, Amnis) was performed on oil-in-water mLDs with 60x magnification. Data was analyzed with the IDEAS software to isolate mLDs of interest through the analysis of the brightfield image via selecting the single (spot count = 1), in-focus (RMS ≥ 55), round (shape ratio ≥ 0.5), large (area ≥ 13 μm^2^), and appropriately colored droplets. The imaging flow cytometer enabled fluorescence observation of up to six color channels simultaneously, but only the brightfield images were analyzed here.

### Fluorescence recovery after photobleaching (FRAP)

Fluorescence recovery after photobleaching (FRAP) was performed on an inverted microscope equipped with focused, free-space bleaching lasers set to 1 mW of 488, 561, or 647 nm wavelength and a quad-band dichroic (ZET405/488/561/640m, Chroma) mounted into the top deck of the IX83 (Olympus). Imaging before and after bleaching was performed with widefield LED illumination (SOLIS, Lumencor), fluorescence cubes (49002, 49005, 49006, Chroma), and a sCMOS camera (Zyla, Andor). Typically, images were acquired at 28 Hz and analyzed via the ratio of the intensity in the bleaching spot to the intensity in the surrounding, unbleached monolayer (*I_ratio_*). The normalized ratio of intensities was used to account for LED-caused photobleaching during the acquisition. Analysis was performed by fitting *I_ratio_* to a single exponential according to

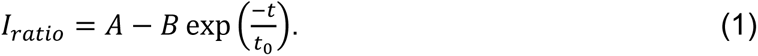

The fitting constants A and B are used to calculate the fraction of the fluorophores bleached by the laser, which is equal to A+B, and the mobile fraction of fluorophores, which equals B/(A+B). The recovery time constant (*t_0_*) and the bleaching width (*w* = 2.5 μm) were used to calculate the observed diffusion coefficient (*D*) according to *D* = *w*^2^/2/*t_0_*.

### Fluorescence correlation spectroscopy (FCS)

FCS was performed with an inverted microscope (IX71, Olympus) that was customized to resolve up to four fluorophores as described previously.^26^ The optical setup included a super-continuum laser (SC-Pro, YSL photonics) that was separated into up to four narrow-spectrum excitation channels by dichroic mirrors and excitation filters (BrightLine FF02-435/40, FF01-513/13, FF01-561/14, and FF01-630/38, Semrock). Each excitation channel passed through custom beam-expanding telescopes and neutral density filters prior to being recombined. The combined excitation colors were reflected off three-band dichroic mirror (ZET442/514/561m, Chroma) to the 40x, 1.30NA microscope objective (UIS2 BFP1, Olympus) to illuminate a diffraction-limited volume within the sample. The emission was transmitted through an emission filter (ZT442/514/561rpc, Chroma), a confocal aperture (P40D, ThorLabs), and relay lenses (Optomask, Andor) that were customized with the insertion of a chromatically separating prism (PS812-A, ThorLabs). The chromatically spread emission was collected on the center pixels of an EMCCD (iXon-897 Ultra, Andor) controlled via custom LabVIEW routines (National Instruments). Samples were exposed to <10 μW total excitation of wavelengths 450, 515, and 561 nm. The far-red fluorophores (*i.e.*, Cy5) were sufficiently excited by the 561 nm light such that no power was needed by the 635 nm excitation channel. Each camera frame was acquired with a 1 ms exposure time that gave 805 Hz frame rate frame rate that yielded 24000 frames per 30 sec acquisition for the 496x4 cropped region of interest on camera. We recently demonstrated multi-color fluorescence cross-correlation spectroscopy (FCCS).^26^ In this manuscript, we focus on concentration-dependent, single-color FCS.

For each time point, each camera acquisition was averaged over the vertical dimension and analyzed as an intensity versus wavelength along the horizontal dimension, which was mapped to the emission color. The control samples of individual fluorophores gave fluorescence emission calibration spectrum which were shifted for sub-pixel alignment and linearly fit to the multi-fluorophore emission spectra of each frame.^26^ The intensity of each fluorophore versus time (*I*(*t*)) was extracted. The autocorrelation (*G*) of a single *I*(*t*) reveals its diffusive behavior according to

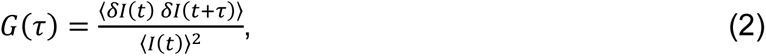

〈*I*(*t*)〉 represents the time average intensity, and δ*I*(*t*) = *I*(*t*) − 〈*I*(*t*)〉. The autocorrelations were fit to the expected form of 2D Brownian diffusion to extract the correlation amplitude (*G_0_*) and the characteristic diffusive dwell time (τ_*D*_) according to

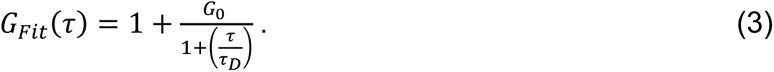

The diffusers diffusion coefficient (*D*) is calculated according to 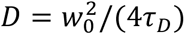 with the width of the observation spot (*w_0_* = 210 nm).

### Fluorescence lifetime imaging microscopy (FLIM)

Fluorescence lifetimes of Flipper-TR (Cytoskeleton, Inc.) were acquired on a MicroTime 200 (PicoQuant) equipped with an inverted confocal microscope (IX73, Olympus) and a 75 µm pinhole. Samples were excited with a 485 nm wavelength pulsed diode laser (LDH-D-C-485 PicoQuant) at a repetition rate of 20 MHz. Excitation light was delivered, and fluorescence was collected by a 100x oil-immersion objective lens (UPLSAPO100XO, Olympus). The fluorescence emission was collected on a single-photon avalanche diode (Photon Counting Module, Excelitas) after blocking the laser excitation with a dichroic mirror (ZT473/594rpc, Chroma) and a long-pass filter (488LP, Semrock). The photon emission delay times (*t_delay_*) relative to the laser pulse were recorded with TCSPC timing electronics (HydraHarp400, PicoQuant) in T3 mode. The delay time histogram (*H*) was fit with a biexponential providing two decay times (*t_1_* and *t_2_*) according to

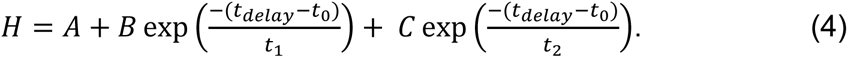

The meaningful fluorophore emission time was the longer of *t_2_* and *t_1,_*^28^ which we indicate as *τ*.

FLIM images of mLD caps were created by raster scanning over the region of interest with 0.2 µW average power and 1 ms dwell time per pixel. The photon delay times in each pixel were fit to Eq. 4 and displayed as the pixel hue. The total number of photons collected per pixel was displayed as pixel brightness. Alternatively, randomly diffusing small, buoyant mLDs were analyzed by focusing 10 μW of average laser power into at 15 μL aliquot on a coverslip and collecting bursts of emission until the delay time histogram was well resolved.

Analysis was done with a custom Python script that selected only the bursts of photons above background. The non-linear fitting of Eq. 4 and burst identification was performed via multiple methods to identify the robustness of the fitting results and the correlation between fit parameters. The background intensity (*A*) was set equal to the mean of *H* for *t* > 35 ns. Fitting for the four unknowns of *B*, *C*, *t_1_*, and *t_2_* yielded more sample-to-sample variations for *τ* than keeping *t_1_* fixed and fitting for only *B*, *C*, and *t_2_*, assuming *t_2_* > *t_1_*. We performed a wide variety of fitting constraints to identify the values of *τ* robust to our manually set fitting parameters. Many fits to each data set were performed, including *t_1_* set equal to a value between 1.25 to 4 ns while keeping only bursts of emission that exceeded 4 to 12 kcounts/sec, only fitting only *t* ≥ 1.5 to 6 ns, requiring a maximum value of *H* ≥ 400 counts within a 16 ps bin, and only keeping *τ* values that were fit with an uncertainty less than 0.15 ns.

### Atomic force microscopy (AFM)

Atomic force microscopy (AFM) was performed with a Bioscope Catalyst (Bruker) mounted upon an inverted optical microscope (IX81, Olympus). The AFM was equipped with colloidal tip cantilevers that provided a spherical tip of 5-μm radius, and a spring constant of 0.1 N/m (CP-qp-CONT-BSG, Nanoandmore). The spring constant for each tip was measured via the thermal tuning method. The tips were cleaned with Milli-Q water and isopropanol prior to each measurement. The colloidal AFM tip was optically aligned with the center of the mLD cap and approached with a speed of 100 nm/s until a repulsion force of 300 pN was measured, after which the tip was retracted at 100 nm/s. All force curves were assessed to ensure no force was observed at the greatest retraction distances, consistent with no contact between the mLD cap and the cantilever.

The force applied to the cantilever tip was linearly fit for repulsion forces between 100 and 300 nN, which typically occurred within 100 nm of first contact with the >30 μm wide mLD caps. The slope of the force to tip height was converted to surface tension by assuming the mLD cap deformed from a hemisphere towards a cylinder, which is represented as

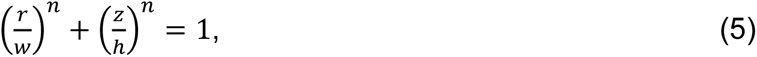

The bud was modeled in cylindrical coordinates with azimuthal symmetry by the radius (*r*), axial coordinate (*z*), a contact radius on the coverslip (*w)*, a height above the coverslip (*h*), and the transition between a sphere and cylinder by the variable *n*. Prior to AFM compression, we assume the mLD cap was a hemisphere with *n =* 2 and *h* = *w.* As the AFM cantilever pressed upon the mLD cap, *h* decreased and *n* increased with a constant droplet volume and glass contact area. We numerically solved for the increase in *n* and the mLD cap area (*A*) with decreasing *h* (Fig. S5); our custom Python script is freely available for download.^34^ The approximately linear relationship between force to depress the mLD cap (*F*), the cap surface tension (*γ*), and the change in *A* versus *h* enabled our fitting of the directly measured *F* versus *h* to determine *γ* according to

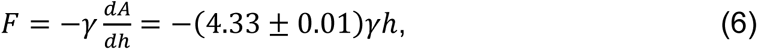

Measurements were repeated with 20 force curves on each of three mLD caps per sample on three separate samples per condition. The force curves that demonstrated an adhesion to the mLD ≥ 10 pN were excluded from analysis. The ramp distance was set to 2500 nm to ensure that a plateau of no force was observed when the tip was far from the mLD, and the approach was halted with a peak-force mode set to 300 pN to avoid excessive compression of the mLD or submersion of the colloidal tip within the TAG.

### Pendant droplet tensiometer

A custom pendant droplet tensiometer was created to measure the surface tension of the TAG-buffer interface upon the addition of PLs. The droplet volume was controlled by a syringe pump (NanoJet, Chemyx) with a 1 mL TAG-filled syringe (1001 LT SYR, Hamilton) connected via a PEEK tubing to a 30-gauge blunt-tipped needle (930050-90BTE, Metcal) within a polystyrene cuvette (759076D, BrandTech). The droplet was imaged with CMOS camera (MU233-FL, AmScope), tube lens (TTL180-A, ThorLabs), and 4x air objective (PlanN, Olympus) after illumination with a diffused (DG10-1500, Thorlabs) desktop LED lamp (DLST01-S, Newhouse). Images were analyzed and the surface tension was extracted with FIJI and OpenDrop.^31, 32^ FIJI was used to convert the

## Acronyms

AFM: – atomic force microscopy
CFP: – cyan fluorescent protein
Cy5: – 1,2-dioleoyl-sn-glycero-3-phosphoethanolamine-N-Cyanine 5
DEV: – droplet embedded vesicle
ER: – endoplasmic reticulum
FCS: – fluorescence correlation spectroscopy
FLIM: – fluorescence lifetime imaging microscopy
FRAP: – fluorescence recovery after photobleaching
GUV: – giant unilamellar vesicle
IB: – intracellular buffer
LD: – lipid droplet
LUV: – large unilamellar vesicle
mLD: – model lipid droplet
MLV: – multilamellar vesicle
PC: – 1,2-dioleoyl-sn-glycero-3-phosphocholine
PE: – 1,2-dioleoyl-sn-glycero-3-phosphoethanolamine
PI: – 1,2-dioleoyl-sn-glycero-3-phosphoethanolamine
PL: – phospholipid
PLIN: – perilipin
TAG: – triacylglycerol
TF: – 2-distearyl-sn-glycero-3-phosphoethanolamine-N-(TopFluor AF488)
YFP: – yellow fluorescent protein

## Supplemental Information

The supplemental materials include seven figures: Figure S1. Z-stack of mLDs; Figure S2. TAG fragmentation; Figure S3. Confocal reconstruction of mLD caps; Figure S4. FRAP images; Figure S5. Numerical cap shape determination; and Figure S6. Pendant droplet volume versus time.

## Acknowledgements

This material is based upon work supported by the National Science Foundation under Grants No. DMR-1652316 (C.V.K.) and DMR-1229284 (P.M.H.). Research reported in this publication was supported by the National Institute of Diabetes and Digestive and Kidney Diseases of the National Institutes of Health under award number R01-DK076629 (J.G.G. and C.V.K.). The work was supported in part by a Defense Threat Research Agency CB Technologies Service Academy Research Initiative Grant (K.T.). The Microscopy, Imaging and Cytometry Resources Core at WSU is supported, in part, by the National Institutes of Health under award numbers P30-CA22453 and R50-CA251068-01. We are grateful to the Richard Barber Interdisciplinary Research Program for financial support. Figures 1A-D, 2A, and 3A were created with BioRender.com.

## Author Contributions

All authors contributed to writing the manuscript. C.V.K contributed to the design and analysis of all experiments and simulations. S.A.G contributed to the design and analysis and performance of the buoyant mLD and FCS experiments. C.V.K., N.J., and M.S. performed the GUV and DEV experiments. S.P. and N.J. performed the mLD cap fluorescence experiments. C.V.K. performed the flow cytometry experiments. R.T. performed the AFM experiments. R.T. and P.M.F. contributed to the design and analysis of the AFM experiments. M.A. optimized and performed the pendant droplet experiments. K.T. performed the FLIM experiments and contributed to their design and analysis. M.A.S. and J.G.G. optimized and performed protein expression and purification experiments in addition to contributing to the design of all fluorescence experiments.

## Supplemental Figures

**Figure S1.**
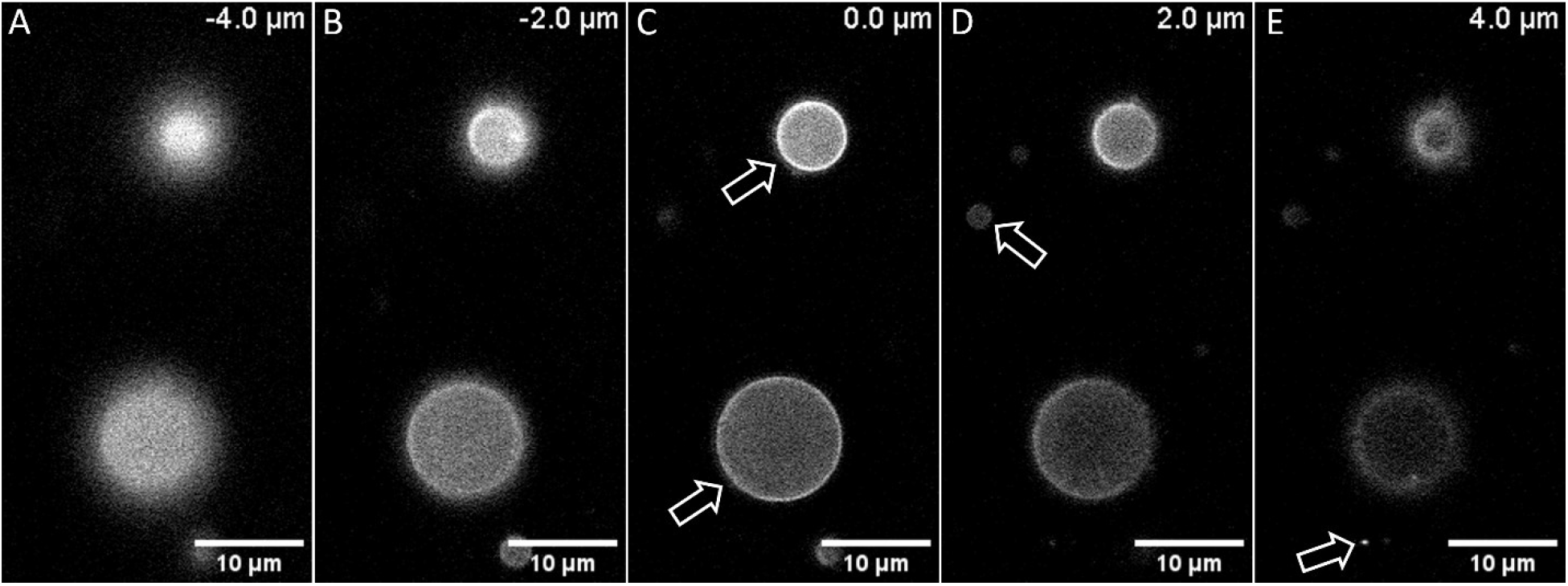
Z-stack of mLDs. (A-E) Widefield fluorescent images from an inverted microscope of buoyant mLDs within a glass imaging chamber with increasing focal planes, starting (A) 4 µm below the equatorial plane of the largest mLDs and ending (E) at top glass slide. (C) The largest mLDs, indicated by the two arrows, have the sharpest contrast at this focal plane. (D) Focusing higher shows equatorial plane of a smaller mLD (arrow). (E) Focusing on the top glass slide shows the smallest mLDs in focus (*arrow*).

**Figure S2.**
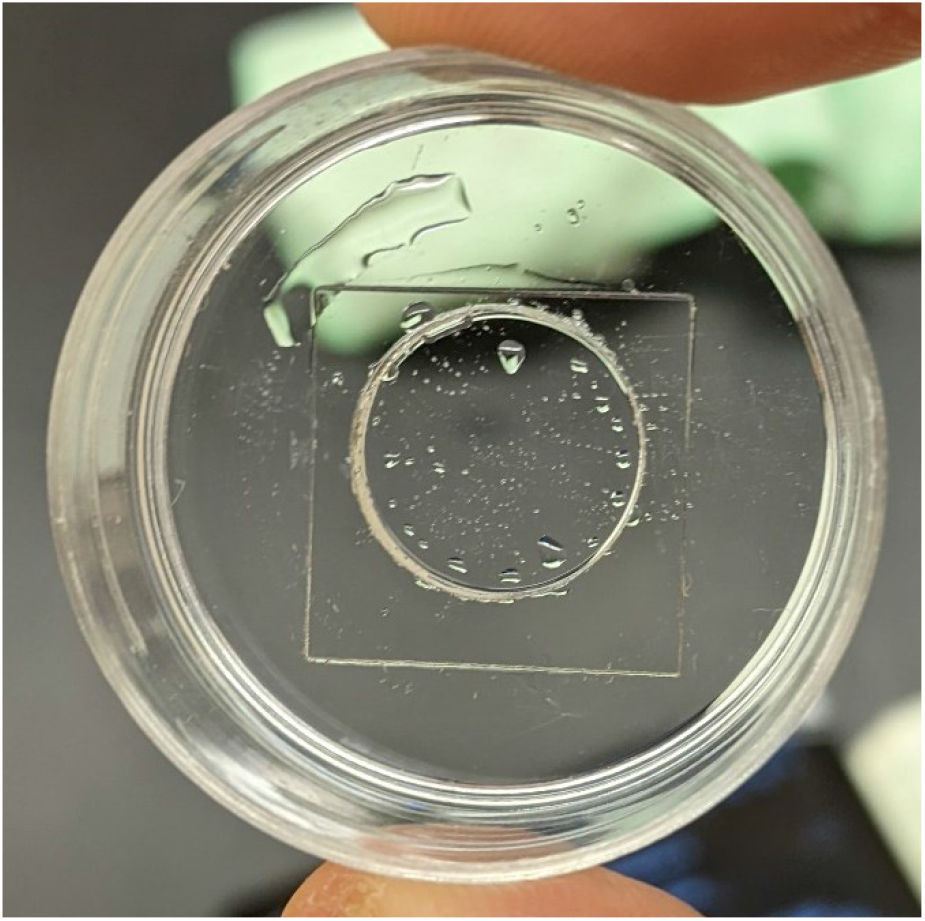
TAG fragmentation. The fragmentation of TAG by an N_2_ stream on a 35-mm diameter glass-bottom dish results in many small oil droplets scattered across the coverslip. These droplets are sufficiently stable and bound to the coverslip such that aqueous buffer may be added to the dish and exchanged without excessively disturbing the <50-μm diameter droplets.

**Figure S3.**
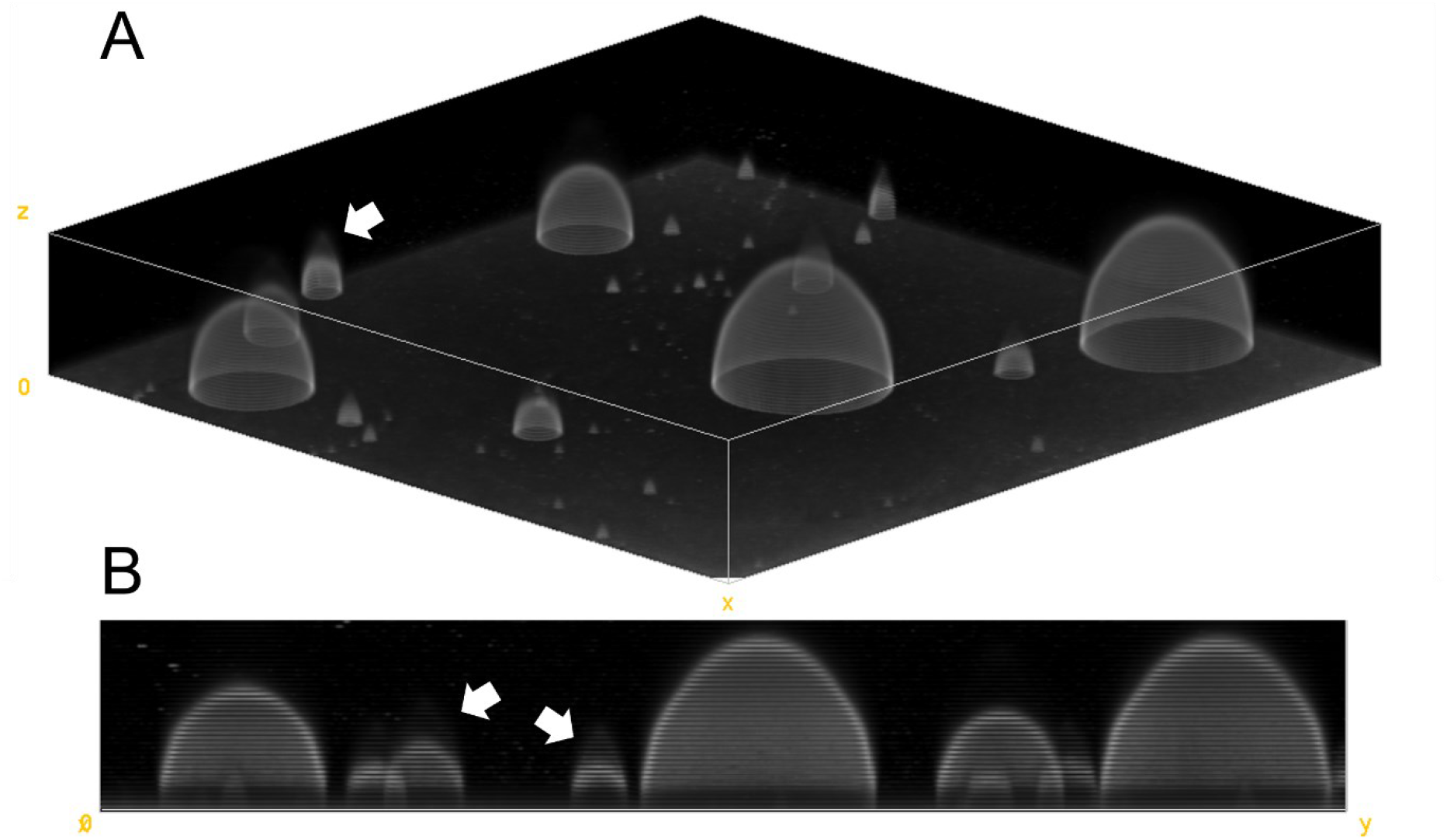
Confocal reconstruction of mLD caps. (A and B) 3D views of confocal images of mLD caps coated with PLIN5-YFP. The view from an angle or (B) parallel to the x-axis shows the near hemispherical shape of the caps. Optical distortions occur when viewing through the curved mLDs results in artifacts above the droplets (*white arrows*). These images represent an extent of 196 μm in each of the x- and y-axes and an extent of 20 μm in the z- axis.

**Figure S4.**
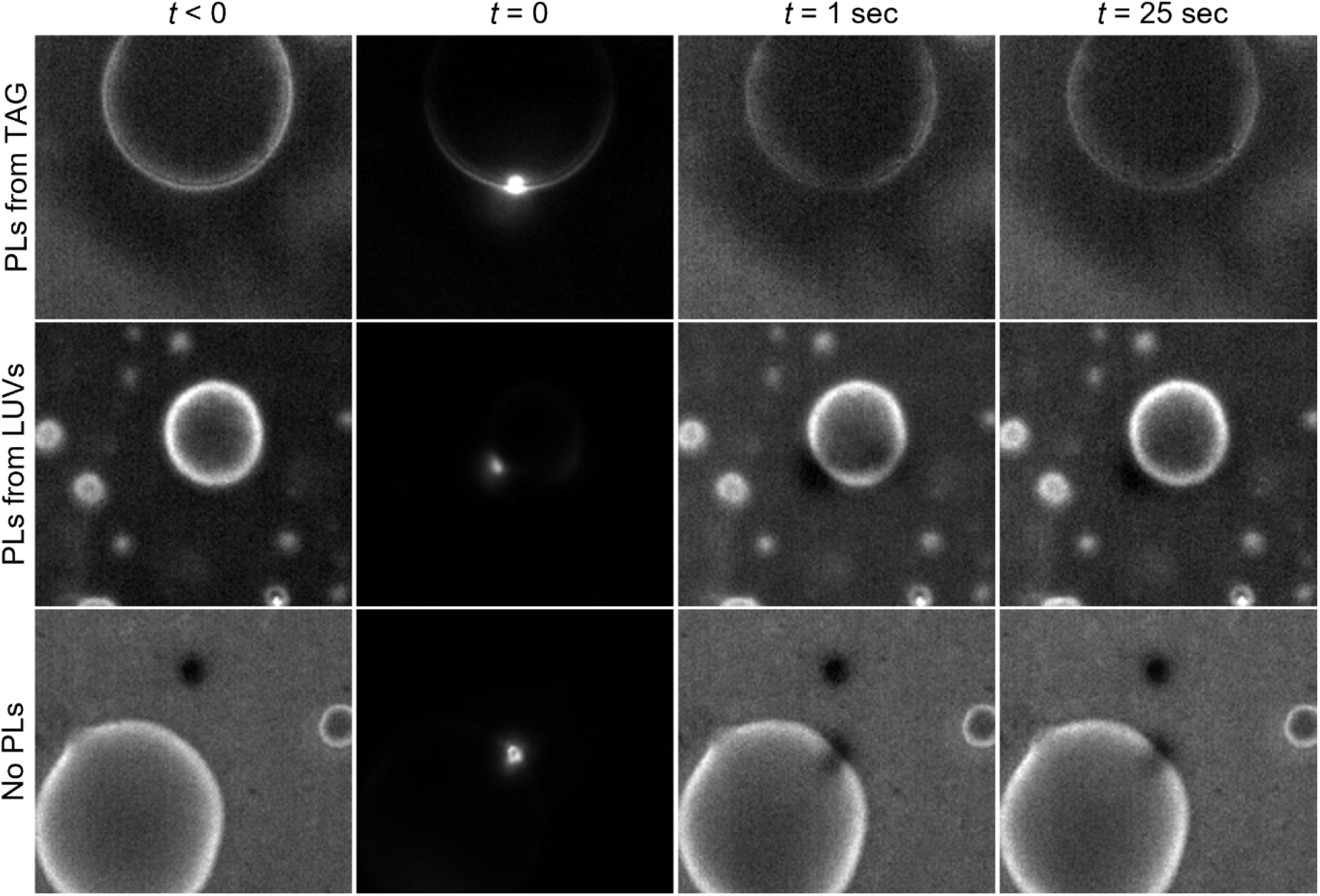
FRAP images. Images taken during FRAP analysis of PLIN5-YFP on mLD caps. The PL monolayer, if present, was either created by PL initially dissolved within the TAG or by exposing the TAG to LUVs. The location of the focused bleaching laser is shown at time (*t*) equal to 0. Note PLIN5-YFP bound to the glass in addition to the mLD; the glass-bound PLIN5-YFP signal was subtracted from the image analysis to obtain the recovery versus time (Fig. 5B). All images represent a width of 16.3 μm.

**Figure S5.**
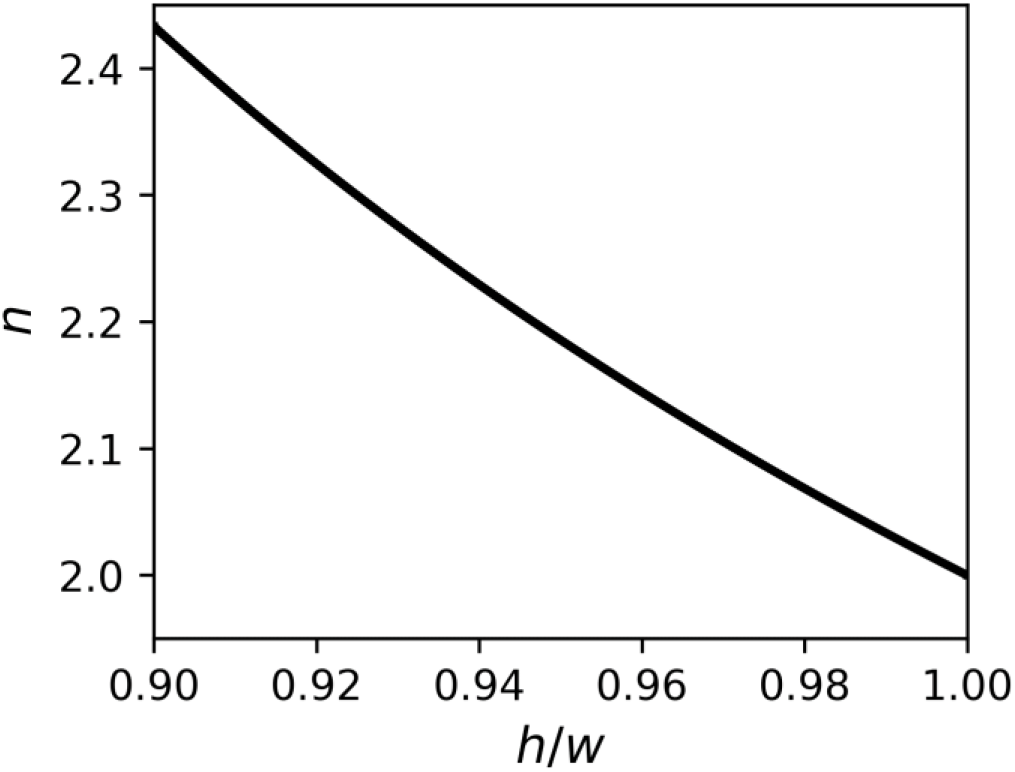
Numerical cap shape determination. The numerically determined relationship between the exponential value *n* (Eq. 5) to model caps of varying height with constant width and volume. The custom Python script used to generate these data is freely available for download.^34^

**Figure S6.**
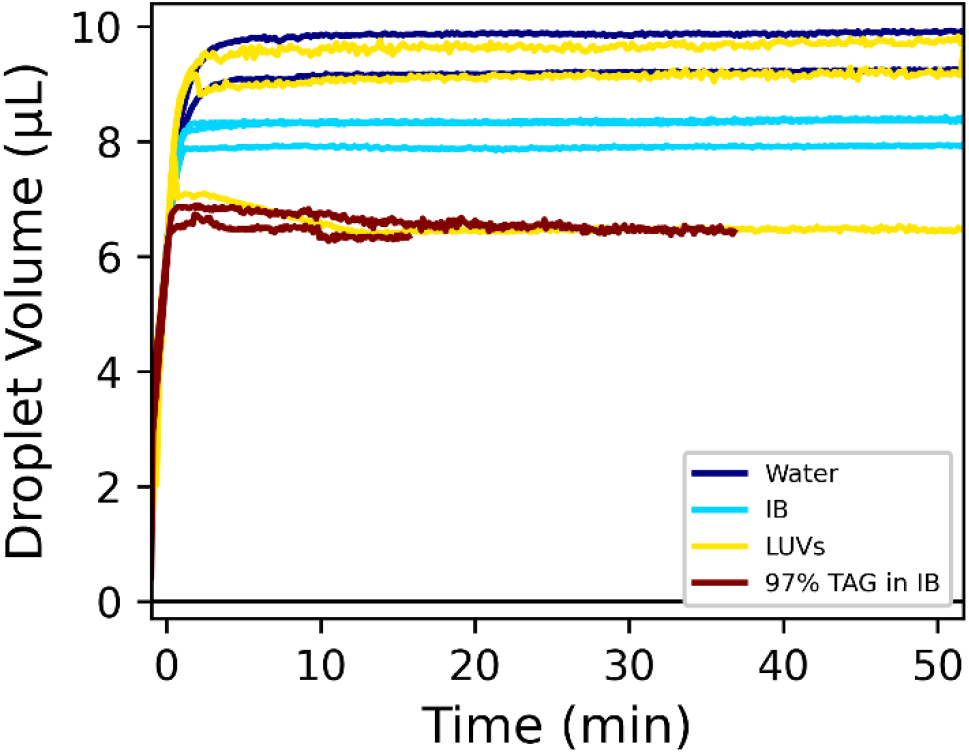
Pendant droplet volume versus time. The syringe-pump controlled droplet size shows a consistent droplet growth rate. Upon stopping the syringe pump at a consistent droplet volume of 6 μL, sample-to-sample variation resulted in the droplet continuing to grow. However, no correlation was observed between the droplet volume and tension.

